# Integrated Low-Energy and Low Carbon Shortcut Nitrogen removal with Biological Phosphorus Removal for Sustainable Mainstream Wastewater Treatment

**DOI:** 10.1101/772004

**Authors:** Paul Roots, Fabrizio Sabba, Alex F. Rosenthal, Yubo Wang, Quan Yuan, Leiv Rieger, Fenghua Yang, Joseph A. Kozak, Heng Zhang, George F. Wells

## Abstract

While enhanced biological phosphorus removal (EBPR) is widely utilized for phosphorus (P) removal from wastewater, understanding of efficient process alternatives that allow combined biological P removal and shortcut nitrogen (N) removal, such as nitritation-denitritation, is limited. Here, we demonstrate efficient and reliable combined total N, P, and chemical oxygen demand removal (70%, 83%, and 81%, respectively) in a sequencing batch reactor (SBR) treating real mainstream wastewater (primary effluent) at 20°C. Anaerobic – aerobic cycling (with intermittent oxic/anoxic periods during aeration) was used to achieve consistent removal rates, nitrite oxidizing organism (NOO) suppression, and high effluent quality. Importantly, high resolution process monitoring coupled to *ex situ* batch activity assays demonstrated that robust biological P removal was coupled to energy and carbon efficient nitritation-denitritation, not simultaneous nitrification-denitrification, for the last >400 days of 531 total days of operation. Nitrous oxide emissions of 2.2% relative to the influent TKN (or 5.2% relative to total inorganic nitrogen removal) were similar to those measured in other shortcut N bioprocesses. No exogenous chemicals were needed to achieve consistent process stability and high removal rates in the face of frequent wet weather flows and highly variable influent concentrations. Process modeling reproduced the performance observed in the SBR and confirmed that nitrite drawdown via denitritation contributed to suppression of NOO activity.

## 1. Introduction

Nitrogen (N) and phosphorus (P) are key limiting nutrients in surface waters, and their removal from wastewater is becoming increasingly important due to widespread eutrophication in both marine and lacustrine environments. While denitrification with exogenous carbon addition to remove N as well as chemical precipitation to remove P are well-established methods to meet nutrient discharge limits, utilities are seeking more efficient and cost-effective methods to meet their permits. Enhanced biological P removal (EBPR) is increasingly implemented as an economical alternative to chemical P precipitation, and emerging innovations in shortcut N removal processes, including nitritation coupled to heterotrophic denitritation via out-competition of nitrite oxidizing organisms (NOO) (Corominas et al., 2010), offer a route to low-energy, low-carbon biological N removal ^2^. However, the drivers that select for NOO out-competition in shortcut N removal processes and their impact on biological P removal are little understood.

While several studies have proposed 2-stage systems with separate sludge for N and P removal ^3–5^, single sludge systems simplify operations and maintenance and can reduce both capital and ongoing costs over 2-stage systems. A limited number of lab-scale studies have used single-sludge systems to incorporate shortcut N removal with P removal from synthetic wastewater feed (Lee et al., 2001; Tsuneda et al., 2006; and Zeng et al., 2003a). Given that chemical oxygen demand (COD) can be limiting in nutrient removal systems, it is important to note that all three of the referenced studies used readily biodegradable acetate in the synthetic feed as their primary carbon source in 10:1 g acetate-COD:gN and 27:1 g acetate-COD:gP ratios or higher. While promising proof of concepts, use of synthetic feed at such high VFA:N and VFA:P ratios is not representative of the dynamics in N, P, and COD composition commonly found in real wastewater.

Investigations of combined shortcut N and P removal from real wastewater without exogenous carbon or chemical addition for P precipitation are limited to one lab-scale reactor ^9^ and two full scale processes ^10,11^, but all three had average wastewater temperatures between 26 and 30 °C. Such elevated temperatures confer a significant advantage to ammonia oxidizing organisms (AOO, which can include both ammonia oxidizing bacteria and archaea) over NOO, thereby greatly facilitating NOO out-competition ^12^, but are not representative of conditions found in WWTPs in temperate regions. In the lab-scale reactor cited above, for instance, Zeng et al. (2014) ^9^ lost NOO out-selection when the wastewater temperature dropped below 23 °C as winter approached. Research into combined shortcut N and EBPR processes with real wastewater at moderate temperatures (i.e. ≤ 20 °C), where NOO suppression is significantly more challenging ^13^, is currently lacking. Intermittent aeration is one promising strategy for NOO suppression at moderate temperatures. Explanations for its efficacy range from a metabolic lag phase of *Nitrospira* NOO compared to AOO upon exposure to oxygen ^14^ to transient exposure to free ammonia due to pH shifts in biofilms ^15^, as free ammonia has a greater inhibitory effect on NOO than AOO ^16,17^. However, the mechanism and efficacy of intermittent aeration for NOO suppression at moderate temperatures, with or without integration of biological P removal, is currently not well understood.

The propensity for shortcut N removal systems to produce nitrous oxide (N_2_O), a potent greenhouse gas, is little understood, though reports suggest that N_2_O production may exceed that of conventional N removal biotechnologies ^18–21^. For example, in Zeng et al., (2003a), N_2_O production exceeded N_2_ production from a lab-scale nitritation-denitritation process by more than 3-fold. However, none of the above studies using real wastewater ^9–11^ measured N_2_O emissions. Therefore, N_2_O measurements on shortcut N removal systems integrated with biological P removal from real wastewater are of interest to accurately assess their net impact on greenhouse gas emissions.

Here, we demonstrate efficient and reliable combined shortcut N, P, and COD removal in a sequencing batch reactor (SBR) treating real mainstream wastewater (primary effluent) at 20°C. In contrast to the synthetic studies cited above, the primary effluent used here as influent contained average ratios of 1:1 gVFA-COD:gTKN and 8.2:1 gVFA-COD:gTP, comprising a challenging environment for total nutrient removal. Importantly, EBPR was coupled to nitritation-denitritation for energy and carbon-efficient N removal. A simple kinetic explanation for the out-competition of NOO via intermittent aeration and SRT control was illustrated via batch tests and process modeling. No exogenous chemicals were needed to achieve consistent process stability and high removal rates in the face of frequent rain events and highly variable influent concentrations.

## 2. Materials and Methods

### 2.1 Reactor inoculation and operation

A 56-L reactor was seeded with activated sludge biomass from another pilot EBPR bioreactor (grown on the same wastewater) on June 15, 2017 (day 0 of reactor operation) and fed primary settling effluent from the Terrance J. O’Brien WRP in Skokie, IL for 531 days. Online sensors included NH_4_^+^, DO, pH and oxidation-reduction potential (ORP) (s∷can, Vienna, Austria). The reactor was operated with code-based Programmable Logic Control (PLC) (Ignition SCADA software by Inductive Automation, Fulsom, CA, USA, and TwinCAT PLC software by Beckhoff, Verl, Germany) as a sequencing batch reactor (SBR) with cycle times detailed in Table 1. An anaerobic react period followed by an intermittently aerated period was chosen with the intent to select for integrated biological P removal and nitritation/denitritation via suppression of NOO activity. The reactor was temperature-controlled to target 20°C (actual temperature = 19.8 ± 1.0°C) via a heat exchange loop to evaluate performance at moderate temperatures. The pH was not controlled and varied between 7.0 and 7.8. NH_4_^+^ sensor-based control was used to control aerobic react length, as detailed in Table 1 and in the Supporting Information. Because react length varied with influent NH_4_^+^ concentration (due to NH_4_^+^ sensor-based control), the SBR loading rate followed that of the full-scale plant, i.e. with shortened SBR cycles and increased flow during wet-weather events. The process timeline is split into 2 phases to simplify reporting: Phase 1 (days 0 −246) and Phase 2 (days 247 – 531), the latter of which represents lower target effluent N concentrations and better N-removal performance. Details on intermittent aeration control can be found in the Supporting Information.

**Table 1.** SBR cycle timing (gravity fill, anaerobic reactor, aerobic react, wasting, settling, and decant) and reactor control details. The end of the SBR aerobic (intermittently aerated) react phase was determined based on an NH_4_^+^ setpoint shown in the table.

|  | *Phase 1* | *Phase 2* |
| --- | --- | --- |
| Days of operation | 0 to 246 | 247 to 531 |
| Gravity fill (min) | 3 to 6 |  |
| Anaerobic react (min) | 45 |  |
| Aerobic react via intermittent aeration (min) | $317 \pm 146$ | $206 \pm 105$ |
| Wasting (min) | 0 to 2.2 |  |
| Settling (min) | 30 to 40 |  |
| Decant of 5/8 volume fraction (min) | 4.5 to 6.0 |  |
| Online $\text{NH}_4^+$-based control target effluent concentration ($\text{mgNH}_4^+\text{-N/L}$) | 3 to 5 | 1.5 to 2 |

SRT was controlled via timed mixed liquor wasting after the aerated react phase, and solids losses in the effluent were included in the dynamic SRT calculation, following the methodology of ^22^. Using an operational definition of “aerobic” as > 0.2 mgO_2_/L, an analysis of 4 cycles from Phase 2 showed that an average 48% of the time within the intermittently aerated react period is aerobic. See the Supporting Information for details regarding SRT control and calculations.

Composite sampling as summarized in Table 2 was initiated on day 27 after an initialization period to allow the accumulation of AOO as measured by ammonia oxidation activity. Beginning on day 114 and to the end of the study, influent COD fractionation analysis was conducted once per week with the following definitions ^23^:

- Particulate COD = Total COD – 1.2-μm filtered COD
- Colloidal COD = 1.2-μm filtered COD – floc-filtered COD
- Soluble COD (not including VFAs) = floc-filtered COD – VFA
- VFA COD = VFA

**Table 2.**
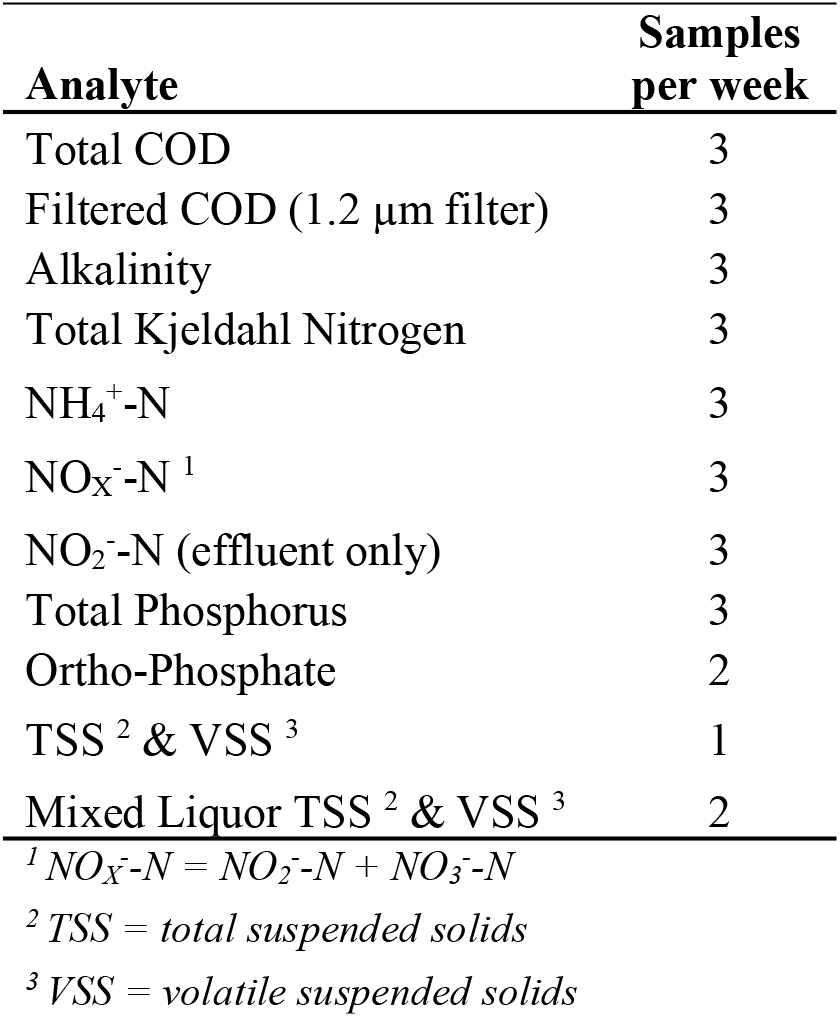
Sampling frequency and target analytes (per APHA, 2005) for daily composite samples. All samples listed are of reactor influent and effluent except NO_2_^−^-N (effluent only) and mixed liquor TSS and VSS (sampled in-reactor).

Floc-filtered COD was measured as described in Mamais et al. (1993) and total COD, filtered COD and VFAs were analyzed per Standard Methods ^25^. On average, the total COD and VFA to nutrient ratios of the influent were (Table S1):

- 8.3:1 g total COD:g TKN
- 1:1 g VFA-COD:g TKN
- 67:1 g totalCOD:g totalP
- 8.2:1 g VFA-COD:g totalP

### 2.2 Batch activity assays

#### 2.2.1 In-cycle batch activity assays

Seventeen in-cycle batch activity assays were conducted throughout the study to monitor *in situ* dynamics of NH_4_^+^, NO_2_^−^, NO_3_^−^, PO_4_^3−^ (all tests), readily biodegradable COD (rbCOD - two tests) and volatile fatty acids (VFAs – one test) via Standard Methods ^25^ and Mamais et al., 1993 for rbCOD. Samples were taken every 15 – 45 minutes for a full SBR cycle, except in the case of two high-frequency tests, in which samples were taken every one to two minutes for 40 minutes in the aerated portion of the cycle to investigate high time resolution nutrient dynamics during intermittent aeration.

#### 2.2.2 *Ex situ* batch activity assays

*Ex situ* maximum batch activity assays for AOO and NOO were performed as previously described ^26,27^. *Ex situ* activity assays were also employed to quantify biological P uptake of polyphosphate accumulating organisms (PAOs) under aerobic and denitrifying conditions. Relative P-uptake rates via different electron acceptors under typical in-reactor conditions was desired (as opposed to maximum P-uptake rates), so external carbon was not added. 250-mL aliquots of mixed liquor were removed from the reactor following the anaerobic phase (i.e. after P release and VFA uptake) and placed in air-tight 250-mL serum bottles. The sealed bottles were injected with sodium nitrite or potassium nitrate stock solutions to approximately 9 mgN/L of NO_2_^−^ or NO_3_^−^ for the anoxic (denitrifying) uptake tests or opened and bubbled with air through an aquarium diffusor stone for aerobic tests. A replicate for the aerobic test was provided by the 56-L reactor itself, which was also aerated continuously (with a resulting DO concentration of 2 mg/L) and sampled in parallel with the aerated serum bottle. A control assay utilized biomass with no electron acceptors (O_2_, NO_3_^−^, or NO_2_^−^) provided. Serum bottles were mixed by a Thermo Scientific MaxQ 2000 shaker table (Waltham, MA) at 150 RPM and at ambient temperature near 20°C. P uptake was quantified via a least squares regression of the PO_4_^3−^ measurement from 3 – 5 samples taken every 20 minutes and normalized to the reactor VSS. The results represent the average ± standard deviation of three total replicates for each electron acceptor from days 237 and 286.

#### 2.2.3 In-cycle batch activity assays for quantification of N_2_O emissions

N_2_O emissions from the reactor were estimated during Phase 2 by measuring the aqueous N_2_O concentration over 8 separate cycles from days 414 to 531 with a Unisense N_2_O Wastewater Sensor (Aarhus, Denmark). N_2_O emissions were calculated from the aqueous concentration following Domingo-Félez et al. (2014), after measuring the N_2_O stripping rate during aeration with mixing and during mixing alone. NH_4_^+^, NO_2_^−^ and NO_3_^−^ were measured concurrently at the beginning and end of cycles ^25^ to calculate TIN removal. N_2_O emissions were then quantified relative to TIN removal and the TKN load for each of the eight cycles.

### 2.3 Process Modeling

To evaluate mechanisms of NOO suppression and the balance between aerobic PAO and denitrifying PAO (DPAO) activity, the SIMBA#3.0.0 wastewater process modeling software (ifak technology + service, Karlsruhe, Germany) was used to simulate performance of the reactor during Phase 2 of operation. The inCTRL activated sludge model (ASM) matrix, based on Barker and Dold (1997) with the addition of two-step nitrification-denitrification, methanotrophs, and other extensions, was utilized without adjustment of kinetic or stoichiometric parameters. Default Monod half-saturation constants of particular relevance to this study include oxygen affinity of AOO (*K*_*02,A00*_ = 0.25 mgO_2_/L) and NOO (*K*_*02,N00*_ = 0.15 mgO_2_/L) and substrate affinity of AOO (*K*_*NHx,A00*_ = 0.7 mgNHX-N/L; NHX = NH_4_^+^ + NH_3_) and NOO (*K*_*N02,N00*_ = 0.1 mgNO_2_^−^-N/L); further parameters can be found in the Supporting Information. SBR control of the reactor was simulated directly using a petri net approach, with sequence control shown as green blocks in Figure S1. To avoid rounding errors and to improve simulation speed, the reactor was modeled with a 56 m^3^ working volume as opposed to 56 L. As in the reactor, the modeled anoxic period was fixed at 45 minutes and the aerobic period ended when soluble NHX (i.e. NH_4_^+^ + NH_3_, which is approximately equal to NH_4_^+^ at the pH values encountered of 7.0 – 7.8) was < 2 mgN/L. Modeled intermittent aeration during the aerobic period was controlled as described in the Supporting Information, though a slightly longer “anoxic” timer of 3 min 45 seconds in the model was used (vs. 0 – 3 minutes in the actual SBR) to account for the DO sensor delay in the actual SBR. Modeled mixed liquor wasting wasted was adjusted until the calculated model SRT (which included effluent solids) matched the SRT of the reactor during Phase 2. 5/8 volume decant was performed at the end of the cycle and average primary effluent (reactor influent) values from Phase 2 were used as model influent. The initialization procedure involved running the model for 150 days to achieve quasi steady-state conditions. Modeled specific growth rates for AOO, NOO, and PAOs were quantified throughout the SBR cycles with rate equations and parameter values from the SIMBA# inCTRL ASM matrix.

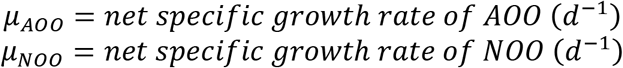

The washout SRT for NOO was calculated from μ_NOO_ as detailed in the Supporting Information.

Modeled PAO growth rates as discussed in this paper include growth on PHA associated with P uptake but do not include decay or PAO growth on PHA where PO_4_^3−^ is limiting. Also, the SIMBA# inCTRL ASM matrix considers only a single PAO population with an anoxic growth factor (*η*_*anox,PA0*_ = 0.33) in the DPAO rate equations to estimate anoxic P uptake (see Supporting Information for full rate equations). The three growth rates below therefore represent growth of a single functional group split between 3 electron acceptors: O_2_, NO_2_^−^, and NO_3_^−^.

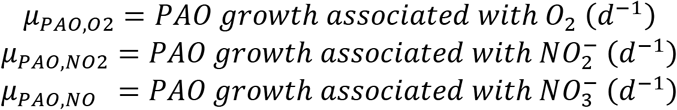

Rate equations and parameters values for the above modeled growth rates, along with the process representation in SIMBA#, can be found in the Supporting Information.

### 2.4 Biomass sampling and DNA extraction

Reactor biomass was archived biweekly for sequencing-based analyses. Six 1 mL aliquots of mixed liquor were centrifuged at 10,000g for 3 minutes, and the supernatant was replaced with 1 mL of tris-EDTA buffer. The biomass pellet was then vortexed and centrifuged at 10,000g for 3 minutes after which the supernatant was removed, leaving only the biomass pellet to be transferred to the −80°C freezer. All samples were kept at −80°C until DNA extraction was performed with the FastDNA SPIN Kit for Soil (MPBio, Santa Ana, CA, USA) per the manufacturer’s instructions.

### 2.5 16S rRNA gene amplicon sequencing

16S rRNA gene amplicon library preparations were performed using a two-step multiplex PCR protocol, as previously described ^29^. All PCR reactions were performed using a Biorad T-100 Thermocycler (Bio-Rad, Hercules, CA). The V4-V5 region of the universal 16S rRNA gene was amplified in duplicate from 20 dates collected over the course of reactor operation using the 515F-Y/926R primer set ^30^. Further details on thermocycling conditions, reagents, and primer sequences can be found in Supporting Information.

All amplicons were sequenced using a MiSeq system (Illumina, San Diego, CA, USA) with Illumina V2 (2×250 paired end) chemistry at the University of Illinois at Chicago DNA Services Facility and deposited in GenBank (accession number for raw data: PRJNA527917). Procedures for sequence analysis and phylogenetic inference can be found in the Supporting Information.

### 2.6 Quantitative Polymerase Chain Reaction (qPCR)

qPCR assays were performed targeting the ammonia oxidizing bacterial *amoA* gene via the *amoA-*1F and *amoA*-2R primer set ^31^, and total bacterial (universal) 16S rRNA genes via the Eub519/Univ907 primer set ^32^. All assays employed thermocycling conditions reported in the reference papers and were performed on a Bio-Rad C1000 CFX96 Real-Time PCR system (Bio-Rad, Hercules, CA, USA). Details on reaction volumes and reagents can be found in the Supporting Information. After each qPCR assay, the specificity of the amplification was verified with melt curve analysis and agarose gel electrophoresis.

## 3. Results and Discussion

### 3.1 Nitrogen, AOO and NOO

#### 3.1.1 Overall Performance and Nitrogen Removal

To demonstrate feasibility and evaluate optimal operational conditions for integrated biological P and shortcut N removal via NOO out-selection at moderate temperatures, we operated lab-scale reactor fed with real primary effluent for 531 days. Reactor operation proceeded in two phases. Reactor performance across both phases is shown in Figure 1 and summarized in Table 3. Phase 1 (days 0-246) established proof-of-concept for the compatibility of N removal via nitritation-denitritation via intermittent aeration with EPBR and allowed for optimization of SRT and the aeration regime (intermittent aeration). P removal was consistent during Phase 1 (average PO_4_^3−^ removal = 83%) excepting aeration failures from reactor control issues around days 80 – 90. Because SRT control was utilized as one of the strategies for NOO out-selection, partial washout of AOO during Phase 1 was occasionally observed when mixed liquor wasting was too aggressive (i.e. total SRT less than 5 days, SRT_AER_ less than 2 days, see Figure S2), resulting in lower NH_4_^+^ oxidation rates and higher effluent NH_4_^+^, after which wasting would be suspended to restore AOO mass. The average TIN removal during Phase 1 was 42% but reached >60% during periods of peak performance. The average TSS during Phase 1 was 1,362 ± 623 mg/L, the VSS was 1,052 ± 489 mg/L, and the HRT was 9.7 ± 3.9 hours not including settling and decant.

**Table 3.**
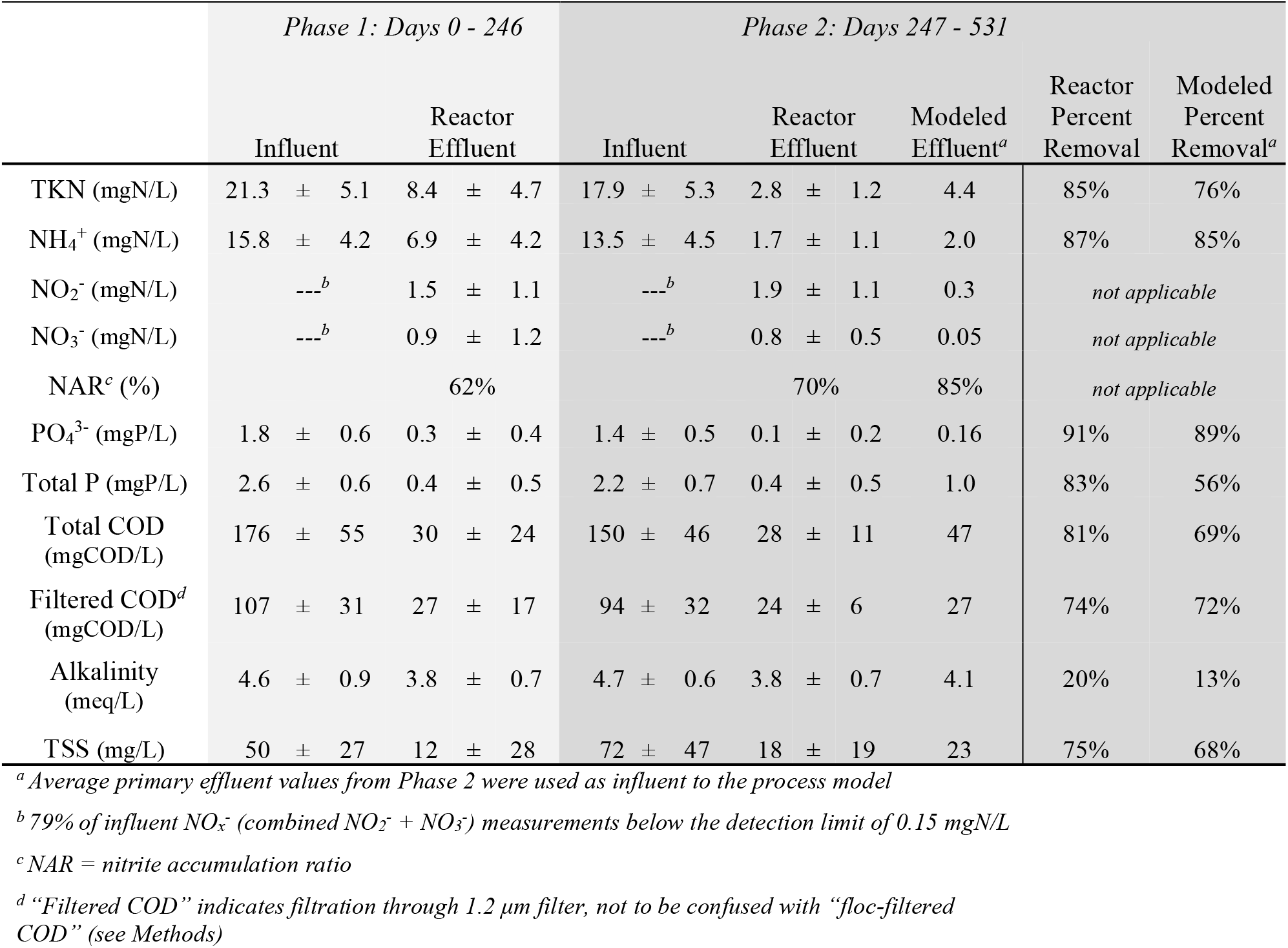
Arithmetic mean ± standard deviation of composite sampling results for influent (primary effluent) and reactor effluent concentrations. Results from Phase 1 are highlighted in light gray and results from Phase 2 are highlighted in dark gray. Process model predictions are for Phase 2 only. Additional information regarding influent COD fractionation can be found in Table S1.

**Figure 1.**
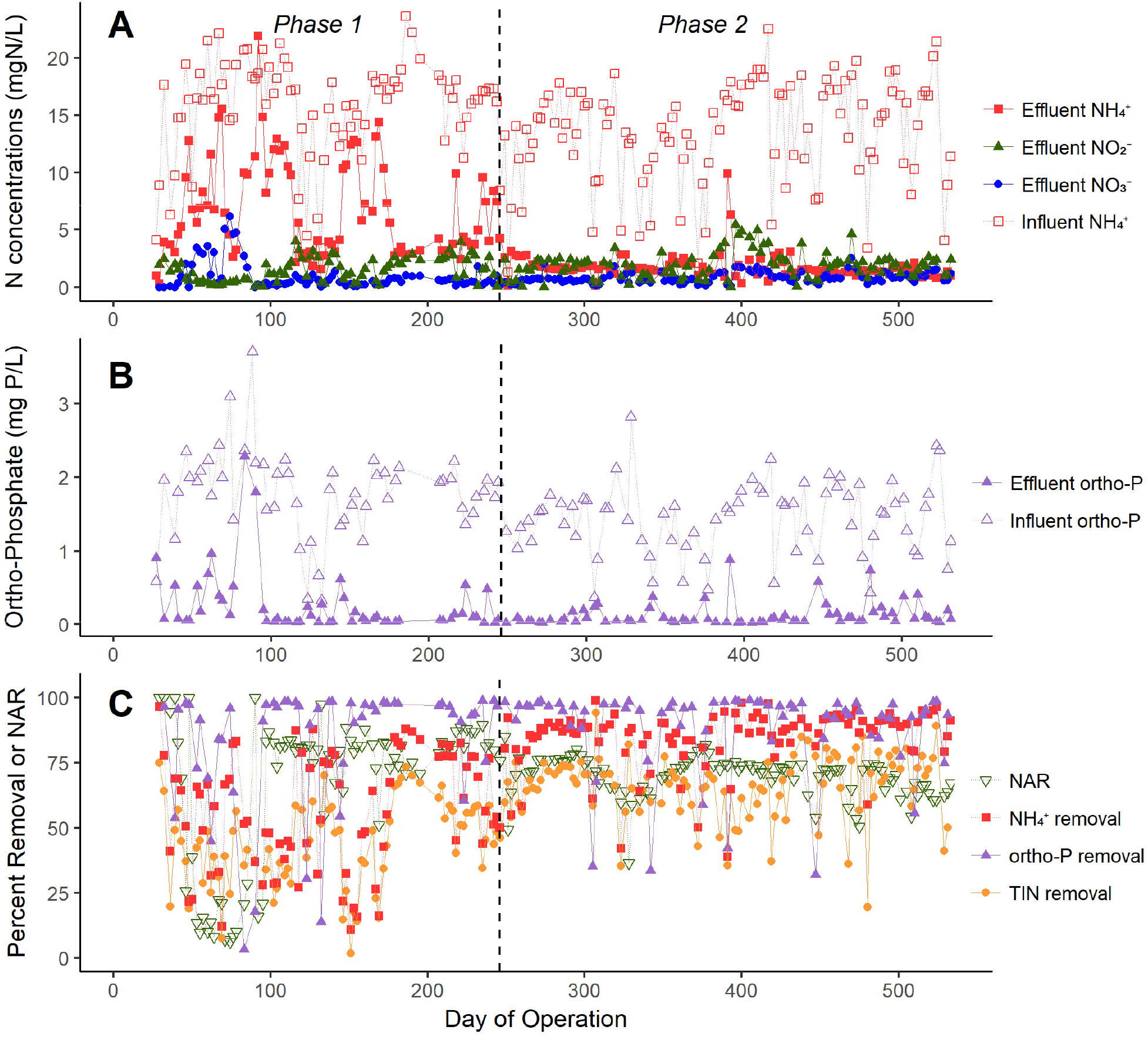
Reactor performance over time from composite sampling (2 – 3 samples/week) over the entire study. **A)** Influent (primary effluent) NH_4_^+^ and effluent NH_4_^+^, NO_2_^−^, and NO_3_^−^. **B)** Influent and effluent orthophosphate. **C)** Nitrite accumulation ratio (NAR) and percent removal of NH_4_^+^, orthophosphate and TIN.

During Phase 2 (days 247-531), SRT control was optimized (total SRT = 9.2 ± 1.8 days, SRT_AER_ = 3.6 ± 0.9 days) and consistent NH_4_^+^ and TKN removal (41 ± 24 mgN/L/d and 54 ± 29 mgN/L/d, respectively, considering influent and effluent values with HRT during Period 2) was achieved while maintaining NOO out-selection (described in section 3.1.2). The average HRT of 6.8 ± 2.8 hours (not including settling and decant) was lower than Phase 1 (9.7 ± 3.9 hours) due to improved AOO activity. Average TIN and PO_4_^3−^ removal during Phase 2 was 68% and 91%, respectively (Table 3). Biological P removal was not impacted by N removal, and the P uptake rate consistently exceeded the NH_4_^+^ removal rate during the aerated portion of the cycle (see Figure 2.A&B for PO_4_^3−^ and NH_4_^+^ concentration profiles through typical cycles). This may have contributed to COD limitation for N removal via denitritation, as COD was most depleted at the end of the SBR cycles (Figure 2.A). This in turn may explain NO_2_^−^ accumulation near the end of most cycles and higher P removal than N removal rates. Figure 2.A and 2.B also demonstrates the variability in react length that was often observed throughout the study due to differences in the NH_4_^+^ oxidation rate, possibly caused by fluctuations in AOO concentrations in the reactor. During Phase 2, the average TSS was 1,773 ± 339 mg/L and the VSS was 1,344 ± 226 mg/L.

**Figure 2.**
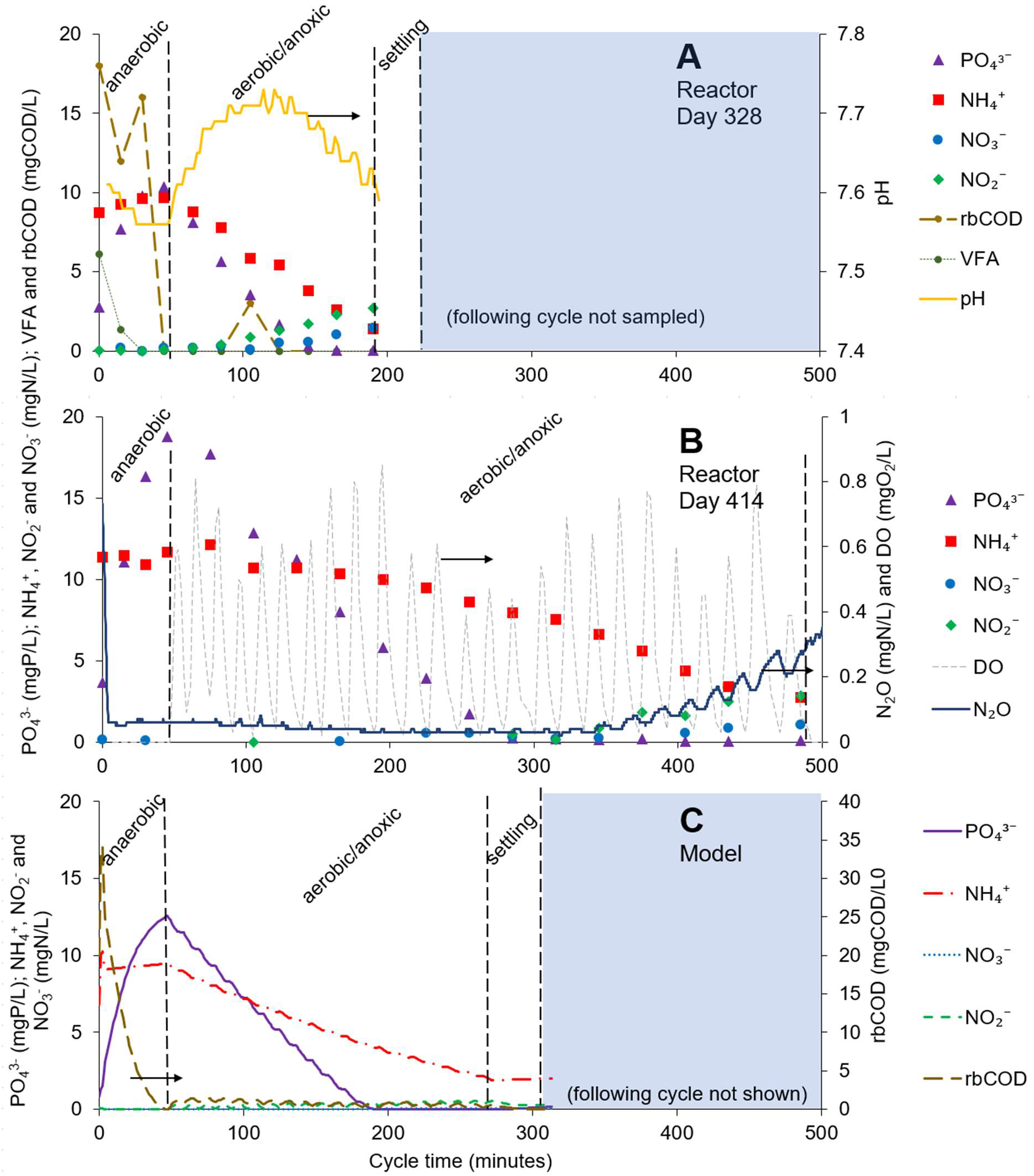
**A) and B)** Two react cycles on days 328 and 414, respectively, that demonstrate efficient P and N removal, selective nitritation, and variability in aerated react length. Cycle **A** included measurements for rbCOD and VFAs, and cycle **B** was run with an N_2_O sensor in the reactor. **C)** SBR cycle as modeled in SIMBA#. rbCOD as shown was calculated as soluble COD*t* – soluble COD_*effluent*_.

#### 3.1.2 NOO Out-selection

A crucial challenge to all shortcut N removal processes, including the nitritation-denitritation with EBPR process that we focus on here, is suppression of NOO activity. To address this challenge, we employed a combination of tight SRT control with intermittent aeration to limit substrate (NO_2_^−^) accumulation. Process monitoring results demonstrated elevated NO_2_^−^ concentrations in the effluent, suggesting successful suppression of NOO activity (Table 3 and Figure 1) with a nitrite accumulation ratio (NAR) of 70% during Phase 2. This observation was corroborated by fifteen in-cycle concentration profiles demonstrating NO_2_^−^ accumulation greater than NO_3_^−^ throughout the cycle (see Figure 2.A&B for two representative cycles). In addition, routine maximum activity assays for AOO and NOO demonstrated that during Phase 2 (optimized, stable reactor operation), maximum AOO activity was 3 to 4-fold greater than NOO (Figure 3).

**Figure 3.**
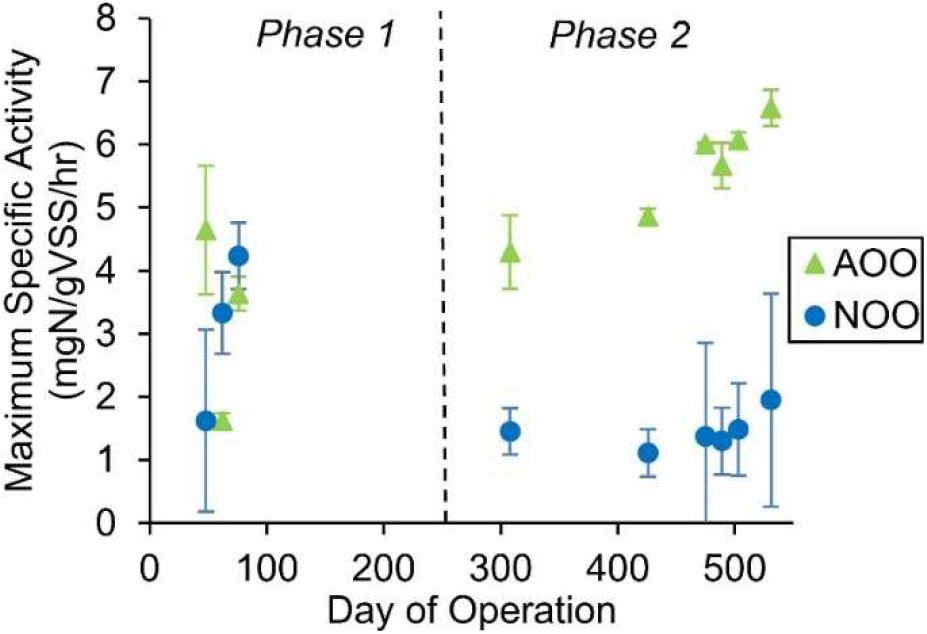
Maximum specific AOO and NOO activity as measured by *ex situ* batch testing. Error bars represent the standard deviation of the method replicates.

To better understand NOO out-selection and nutrient dynamics during intermittent aeration and to provide additional support for suppression of NOO activity in this process, high frequency sampling (1 grab sample/minute for 40 minutes for measurement of NH_4_^+^, NO_2_^−^, NO_3_^−^, and PO_4_^3−^) was conducted during two typical SBR cycles on days 202 and 258 (Figure 4.A, data from day 258 only shown). The resulting concentration profiles show NO_2_^−^ accumulation with very little NO_3_^−^ accumulation during aeration. Two complete intermittent aeration intervals are shown in the early part of the cycle (note that intermittent aeration begins 45 minutes into the cycle), during which NO_2_^−^ accumulates up to 0.4 mgNO_2_^−^-N/L following 5 minutes of aeration, while NO_3_^−^ does not get above 0.1 mgNO_3_-N/L. The NAR during the nitrite peak of these two aeration intervals was 84% and 95%, which demonstrates NOO suppression via selective nitritation. Then, in the subsequent anoxic intervals, the accumulated NO_2_^−^ is drawn down via denitritation. This denitritation provides a robust nitrite sink and one of the methods for NOO out-selection, such that NO_2_^−^ is not available for NOO in the following interval.

**Figure 4.**
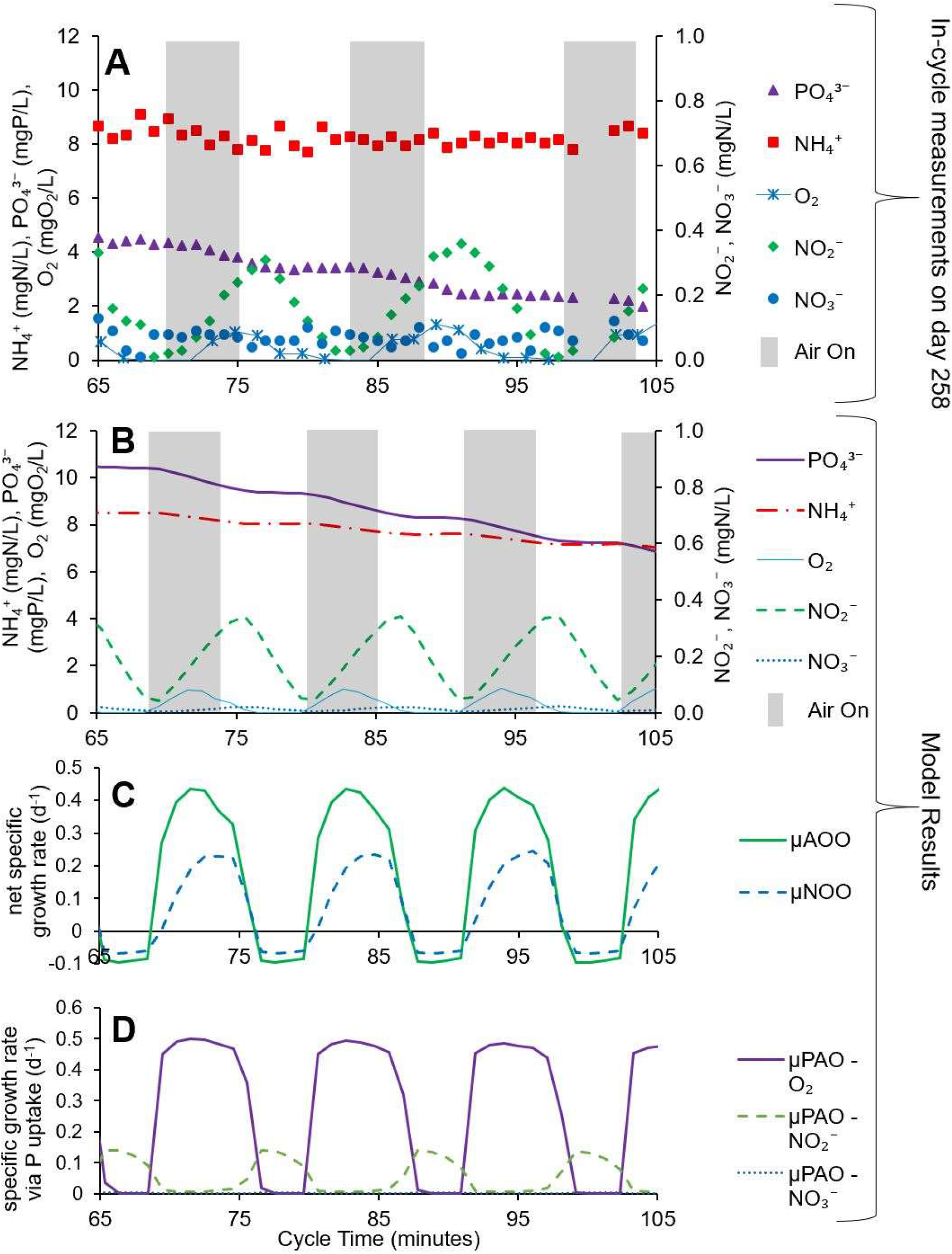
Comparison plot between high resolution within-cycle reactor sampling (A) and modeled results (B, C, D) for the intermittently aerated react period of SBR operation (minutes 65 – 105, beginning 20 minutes after the start of aeration). **A)** Results of grab sampling from a reactor cycle on day 258 of operation. Selective nitritation rather than nitratation during aerated phases (gray shading) is evident and produced NO_2_^−^ is then denitrified in anoxic phases. The s∷can optical DO sensor is rated for a 60-second response time, and a ~1-minute delay is evident in comparison to the model plot B. **B)** Modeled concentration dynamics including on/off switching for aeration control. **C)** Modeled AOO and NOO net specific growth rates including decay. **D)** Modeled PAO specific growth rates associated with P uptake via O_2_, NO_2_^−^ and NO_3_^−^. Decay and growth not associated with P uptake are not included.

Process model results validate the nutrient dynamics observed as seen in Figures 2 and 4. The process model offers additional insight into the mechanism for NOO out-selection. The net specific growth rates of AOO and NOO were calculated from model data output according to rate equations from the inCTRL ASM matrix (see Supporting Information), and are plotted in parallel with the intermittent aeration intervals in Figure 4.C. Due to differences in substrate availability (i.e. high NH_4_^+^ and low NO_2_^−^), μNOO was less than μ_AOO_ at the beginning of each aeration interval and remained below it throughout the 5 minutes of aeration. This specific growth rate differential was maintained throughout much of the cycle, but μ_NOO_ roughly equaled μ_AOO_ by the end of the intermittently aerated react phase due to the accumulation of NO_2_^−^ (data not shown). However, the differential in net specific growth rates in the early part of the SBR cycle ensures that AOO can be maintained in the reactor at a lower SRT than NOO. The modeled average net specific growth rate (including decay) over the cycle can be used to infer a theoretical SRT for NOO to avoid washout, which in this case was 13.2 days (SRT_AER_ = 5.3) days. A similar calculation using the average net specific growth rate of AOO gives an SRT of 8.2 days (SRT_AER_ = 3.3 days), which affirms that AOO are retained via the modeled SRT of 9.5 days. This differential in theoretical SRT (13.2 days for NOO, 8.2 days for AOO) was found with standard kinetic modeling that did not invoke metabolic lag times of NOO (i.e. Gilbert et al., 2014), indicating that substrate limitation alone is sufficient to explain NOO out-competition in this process. The average reactor SRT during Phase 2 was 9.2 ± 1.8 days (SRT_AER_ = 3.6 ± 0.9 days) which, because it is in between the theoretical AOO and NOO SRT values indicated above, reinforces experimental data indicating that SRT control was optimized to washout NOO and retain AOO. Both reactor and modeling results therefore confirm that a combination of intermittent aeration and SRT control can be used to maintain nitritation-denitritation under mainstream conditions. Furthermore, these results suggest that NOO suppression via intermittent aeration and SRT control can be explained by simple substrate (kinetic) limitations alone without invoking more complex mechanisms such as metabolic lag time ^14^ or free ammonia inhibition ^15^.

#### 3.1.3 N_2_O Emissions

N_2_O emissions were measured during 8 separate cycles during steady performance in Phase 2 (between days 414 – 531) and ranged from 0.2 to 6.2% of the influent TKN load, with an average of 2.2 ± 2.0% (Table S2). N_2_O emissions relative to TIN removal averaged 5.2 ± 4.5%. N_2_O accumulation in the reactor generally paralleled NO_2_^−^ accumulation near the end of the aerated portion of the cycle. For example, on the N_2_O test on day 414 (Figure 2.B), grab sampling throughout the cycle revealed that by the time NO_2_^−^ first accumulated above 0.1 mgNO_2_-N/L at 285 minutes, 57% of the TIN removal for that cycle had occurred while only 20% of the N_2_O had been emitted, indicating that relative N_2_O emissions increased in the presence of elevated NO_2_^−^.

The above measurements are comparable to reported N_2_O emission rates for conventional biological nutrient removal (BNR) processes. Ahn et al. (2010) reported a range of 0.01 – 1.8% N_2_O emitted relative to influent TKN at 12 full-scale wastewater treatment plants (WWTPs), which included both conventional BNR and non-BNR processes. Foley et al., 2010 reported a much larger range of 0.6 – 25% N_2_O emitted relative to TIN removed at 7 full-scale conventional BNR WWTPs. Both studies found that N_2_O emissions were correlated with high NO_2_^−^ concentrations, as was the case in our reactor (Figure 2.B). In fact, of the eight cycles analyzed for N_2_O emissions, the four tests with the highest effluent NO_2_^−^ also had the four highest N_2_O emissions. Ahn et al. emphasized that the bulk of N_2_O emissions occur in aerobic zones due to air stripping of N_2_O; indeed, in our reactor 92% of the N_2_O emitted from the in-cycle test on day 414 (for example) occurred during aeration. N_2_O mass transfer (i.e. stripping) coefficients for our reactor were 40 times higher during aeration and mixing than during mixing alone (0.0688 min^−1^ and 0.0017 min^−1^, respectively).

Other shortcut N removal biotechnologies, such as PN/A, have been found to have elevated N_2_O production levels over conventional methods for biological N removal ^18–21^. Both Desloover et al. and Kampschreur et al. (who measured 5.1 – 6.6% and 2.3% N_2_O production relative to influent TKN, respectively) found that a separate nitritation step (as opposed to simultaneous nitritation and anammox) caused increased N_2_O production by AOO, which may be due to elevated NO_2_^−^ concentrations. However, it is not clear that AOO are causing the bulk of N_2_O production in our system or other nitritation-denitritation systems, as low COD concentrations can induce incomplete denitrification and lead to elevated N_2_O production ^35–37^. Indeed, NO_2_^−^ and N_2_O accumulation occurs at the end of the SBR cycles (Figure 2.B) where COD is most depleted from aeration. This suggests that N_2_O emissions from this reactor could be mitigated by a step-feed process, i.e. by filling additional primary effluent to prevent a low COD:N ratio and avoid NO_2_^−^ and N_2_O accumulation at the end of the cycle. Additional research is required to test the effects of this strategy.

An additional potential benefit of a step-feed modification could be a reduction in the effluent NO_2_^−^ concentration. Elevated NO_2_^−^ concentrations in discharge to surface waters is undesirable in part due to its toxicity to fish and other aquatic life ^38^. Aside from a step-feed system, potential solutions to elevated NO_2_^−^ include a final nitrification step (for oxidation of NO_2_^−^ to NO_3_^−^) or an anammox polishing step (as suggested by Regmi et al., 2015). It should be noted that anammox on seeded biocarriers similar to those in the ANITA^TM^Mox process ^40^ could be incorporated into the same reactor for increased N removal, thus eliminating the need for a two-stage system.

### 3.2 P removal and PAOs

Consistent P removal was achieved in Phase 2 and most of Phase 1 (Figure 1, Table 3). EBPR performance was not negatively impacted by long-term nitritation-denitritation; in fact, the P uptake rate exceeded the NH_4_^+^ removal rate throughout the study (see Figure 2.A&B for two representative cycles), indicating that SRT and HRT control to optimize AOO activity (while minimizing NOO activity) ensured sufficient retention and react times for PAOs. The total P removal rate during Phase 2 was 6.8 ± 2.7 mgP/L/d when considering the entire SBR cycle. The P uptake rate from in-cycle testing during Phase 2 was 105 ± 34 mgP/L/d (or 3.4 ± 1.1 mgP/gVSS/hour) when considering the linear portion of P uptake during the aerated react phase (Figure S3).

High frequency sampling (Figure 4.A) and model results (Figure 4.B) both demonstrate P removal during aeration coupled to little to no P removal during periods of anoxia. Importantly, this indicates that P release did not occur in the absence of oxygen, verifying that intermittent aeration with periods of anoxia is compatible with EBPR technologies. However, it also indicates that relatively little denitrifying P uptake occurred, even under anoxic conditions when NO_2_^−^ was present. This suggests that P uptake by aerobic PAO metabolism rather than by denitrifying PAOs (DPAOs) was the predominant driver of P removal. Figure 4.D shows the modeled specific PAO/DPAO growth rates associated with P uptake. Kinetic insights from the process model, which models PAOs as a single group capable of using O_2_, NO_2_^−^ and NO_3_^−^ as electron acceptors for P uptake, show that the combination of low NO_2_^−^ and inhibition due to O_2_ prevented appreciable DPAO activity during intermittent aeration. Modeled P uptake via NO_2_^−^ was only 16% of total P uptake, and modeled P uptake via NO_3_^−^ was even lower at only 0.7% of total P uptake due to limited NO_3_^−^ accumulation. The process model suggests that the presence of residual DO, rather than a lack of NO_2_^−^ or NO_3_^−^, was the primary inhibitor of DPAO activity. Figure 4.D shows that peak DPAO growth in the model occurred not at the maximum NO_2_^−^ concentration (i.e. 75 minutes) but when DO had reached near zero (i.e. 78 minutes), at which point NO_2_^−^ was at about half of the maximum concentration. Finally, while in-reactor, in-cycle measurements of DPAO activity are difficult to make, *ex situ* measurements of P uptake rates via O_2_, NO_2_^−^ and NO_3_^−^ showed that the P uptake via NO_2_^−^ was 17% relative to O_2_, while that of NO_3_^−^ was 14% relative to O_2_ (Figure 5). The high frequency sampling plots, DPAO modeling and *ex situ* P uptake tests all indicate that DPAO activity likely plays a relatively minor role in P removal in this reactor.

**Figure 5.**
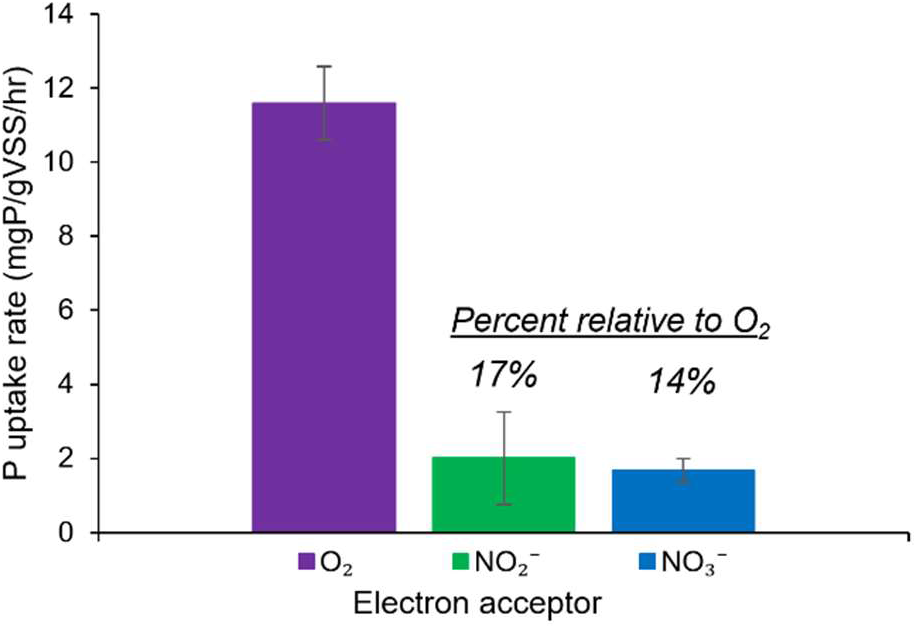
P uptake rates in the presence of O_2_, NO_2_^−^, and NO_3_^−^ from *ex situ* batch tests.

The minor role of DPAOs in this process countered our original expectation that frequent periods of anoxia coupled to the presence of NO_2_^−^ would select for a significant DPAO population. DPAOs are considered advantageous in combined N and P removal processes because they offer the opportunity to reduce carbon demand and aeration requirements ^41^. Lee et al. (2001) were able to achieve 64% DPAO activity (relative to total P uptake) by introducing a single long anoxic phase (with both NO_2_^−^ and NO_3_^−^ present) in the middle of the aerobic phase, which suggests that longer intermittent aeration intervals may select for more DPAO activity (but perhaps at the expense of NOO out-selection). However, preference for DO does not explain the low P uptake via NO_2_^−^ or NO_3_^−^ in the absence of O_2_ (Figure 5) from *ex situ* batch tests in our reactor. Zeng et al. (2003b) observed that *Accumulibacter* PAOs (which were also identified in this study, see Section 3.3) previously acclimated to aerobic P uptake exhibited a 5-hour lag phase in P-uptake when exposed to anoxic conditions (NO_3_^−^) in place of aeration. A metabolic lag phase is unlikely to explain low maximum P uptake via NO_2_^−^ or NO_3_^−^ in this reactor, however, given that linear drawdown of NO_2_^−^ or NO_3_^−^ was observed in all *ex situ* batch tests. A large majority of *Candidatus* Accumulibacter phosphatis genomes sequenced to date have contained the gene encoding nitrite reductase (responsible for reducing NO_2_^−^ to nitric oxide [NO]) ^43^, suggesting that most, if not all, *Accumulibacter* PAOs harbor genomic machinery necessary for denitrifying P uptake via NO_2_^−^. Whether the lack of DPAO activity in this reactor and others is due to the types of PAOs present (and thus the presence or absence of denitrifying genes) or due to the relative expression/inhibition of denitrifying genes present in the PAOs requires further study.

As previously stated, shortcut N removal via nitritation-denitritation did not negatively impact EBPR in this study. Instances of relatively poor P removal were instead usually associated with wet weather flows. Rain not only dilutes the influent but may also induce higher redox conditions in the collection system, indicating a lack of fermentation and little formation of the VFAs that are beneficial to the EBPR process. On sampling days when primary effluent VFAs were at or below the detection limit of 5 mg acetate/L (n = 21), the average PO_4_^3−^ removal of 63% was significantly lower (*p* value = 0.003) than the average PO_4_^3−^ removal of 93% on days when VFAs were greater than 5 mg acetate/L (n = 81).

Shortcut N removal systems can be problematic for EBPR if NO_2_^−^ accumulation leads to elevated concentrations of its conjugate acid, nitrous acid (HNO_2_). HNO_2_ concentrations above 0.5×10^−3^ mgHNO_2_^−^N/L can lead to inhibition of *Candidatus* Accumulibacter PAOs ^45^, which were the dominant PAO identified in this study (see Section 3.3). In the extreme case, the maximum NO_2_^−^ concentration in the effluent of our reactor (e.g. end of the SBR cycle) of 5.4 mgNO_2_^−^-N/L combined with the minimum pH of 7.0 (which did not actually occur simultaneously) corresponds to 0.96×10^−3^ mg HNO_2_^−^N/L with pK_a_ of 3.25 for HNO_2_ 46. This indicates that HNO_2_ was rarely, if ever, above the reported PAO inhibition concentration in our reactor. Moreover, the highest NO_2_^−^ concentrations occurred near the end of the cycle when the majority of PO_4_^3−^ had already accumulated intracellularly as polyphosphate, and residual NO_2_^−^ from the end of the cycle was rapidly depleted after filling at the top of the following cycle.

### 3.3 Functional Guild Analysis: PAO, NOO, and AOO

We used 16S rRNA gene sequencing to evaluate diversity and relative abundance of PAOs, NOO, and AOO in the reactor. *Candidatus* Accumulibacter was the dominant genus of PAO in the SBR throughout the study and ranged in relative abundance from 6.6% to 12.0% (Figure S4). *Tetrasphaera* was detected at most time points but always below 0.3% relative abundance. Glycogen accumulating organisms (GAOs) in the genus *Candidatus* Competibacter, which are potential competitors to PAOs, were consistently less abundant than PAOs, and varied from below the detection limit to 2.4% relative abundance. Other putative GAOs, such as the genera *Defluviicoccus* and *Propionivibrio* ^47^, were found at even lower abundance than *Candidatus* Competibacter (data not shown).

*Nitrotoga* and *Nitrospira* alternately dominated the NOO population according to 16S rRNA gene sequencing (Figure 6). The reason for the alternation is unknown as the timing of succession did not clearly correlate with reactor control or performance, although Keene et al. (2017) observed a similar phenomenon. *Nitrospira* dominated at the beginning of Phase 2, and although the NOO population shifted to *Nitrotoga* over the next 100 – 200 days, there was no corollary change in nitritation-denitritation performance, the NAR, or N removal. This result suggests that the observed robust suppression of NOO activity in this process does not depend upon complete washout of either *Nitrospira* or *Nitrotoga*.

**Figure 6.**
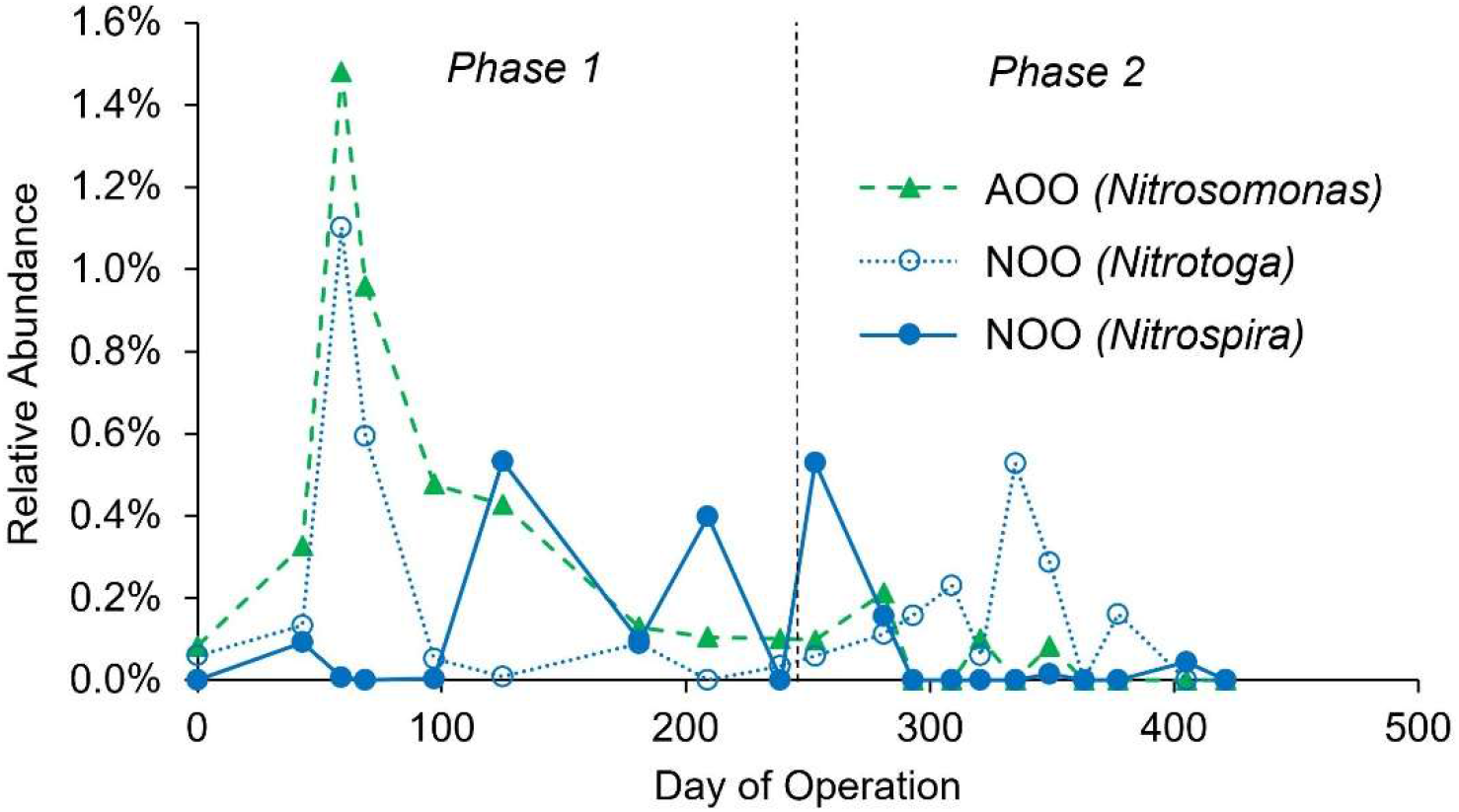
Relative AOO and NOO abundance based on 16S rRNA gene amplicon sequencing through the first 421 days of reactor operation. Day “0” represents the inoculum, which was sampled before reactor operation began.

*Nitrosomonas*-affiliated Betaproteobacteria were the dominant AOO throughout the study according to 16S rRNA gene sequencing but were present at surprisingly low relative abundance for the 2^nd^ half of Phase 1 and all of Phase 2 of reactor operation. Interestingly, the relative abundance of *Nitrosomonas* based on 16S rRNA gene sequencing was below the detection limit for selected samples between days 293 – 431 (Phase 2, Figure 6). No other known AOO were detected during that time; ammonia oxidizing archaea were detected at only two timepoints before day 100 and at low abundance (< 0.04%). Other potential AOO genera, such as *Nitrosospira* and *Nitrosococcus*, were not detected in any 16S rRNA gene sequencing samples. *Nitrospira* can include complete ammonia oxidizing (comammox) clades ^49^, and comammox can in some cases be the dominant AOO ^27^ in wastewater treatment. However, *Nitrospira* were not detected or were at low abundance (< 0.04%) after day 293. The decline in AOO was confirmed by qPCR via the functional bacterial *amoA* gene (Figure S5), although AOO were still detected at all time points via qPCR with a minimum of 0.15% relative abundance on day 421. Although the NH_4_^+^ oxidation rate was variable throughout Phase 2 (Figure S3), NH_4_^+^ oxidation activity was maintained throughout the experimental period. This suggests that either *Nitrosomonas* AOO can maintain effective NH_4_^+^ oxidation rates at very low abundance or an as-yet unidentified organism contributed to NH_4_^+^ oxidation ^50^.

## 4. Conclusions

This study is the first to demonstrate robust combined shortcut N and P removal from real wastewater without exogenous carbon or chemical addition at the moderate average wastewater temperature of 20°C. Mainstream nitritation-denitritation was achieved for more than 400 days via intermittent aeration and SRT control, with an average NAR of 70% during Phase 2. Process modeling reproduced this performance and confirmed that NOO activity was suppressed with a combination of NO_2_^−^ drawdown via denitritation and washout via SRT control, and provided possible explanations for the relative lack of DPAO activity. Importantly, neither NO_2_^−^ accumulation nor periods of anoxia in intermittent aeration adversely affected EBPR performance, and consistent and integrated shortcut TIN and biological P removal were achieved for more than 400 days. N_2_O emissions were in line with observations of other shortcut N removal systems and were primarily associated with NO_2_^−^ accumulation at the end of the cycle. The single-sludge nutrient removal process examined here, as compared to two-stage systems with separate sludges, could reduce operating cost and complexity while meeting nutrient removal goals.

## 6. Acknowledgements

Many thanks to Christian Landis, Adam Bartecki, George Velez, Sandra Matual, Robert Swanson, Thaís Pluth, Thota Reddy, and O’Brien WRP staff and operators.

This study was funded by the Metropolitan Water Reclamation District of Greater Chicago and the National Science Foundation Graduate Research Fellowship under Grant No. DGE-1324585.

