## Supporting Information for "Integrated Low-Energy and Low Carbon Shortcut Nitrogen removal with Biological Phosphorus Removal for Sustainable Mainstream Wastewater Treatment"

### S1. Methods

#### S1.1. Aeration control

The variable-length aerated react period was terminated if either a maximum allowable react time was reached (usually set between 300 – 480 minutes) or if the target $NH_4^+$ concentration (3 – 5 mg$NH_4$-N/L in Phase 1, 2 mg$NH_4$-N/L from days 247 - 430 and 1.5 mg $NH_4$-N/L from days 431 - 531 in Phase 2) was reached according to the online ammo::lyser™ ion-selective electrode (s::can, Vienna, Austria). Intermittent aeration was used during the aerated react period with the following loop:

1. 4 or 5 minutes of aeration with proportional-integral (PI) control to target 1 mg$O_2$/L via the online dissolved oxygen (DO) oxi::lyser™ optical probe (s::can, Vienna, Austria). PI control managed the percent-open time of an air solenoid valve, which, when open, provided compressed air at 7 – 15 liters per minute through a 5-inch diameter aquarium stone disk diffusor at the bottom of the reactor.
2. After aeration, shut air solenoid valve and wait until DO drops to < 0.05 mg$O_2$/L.
3. Run "anoxic" timer for 0 – 3 minutes. At end of timer, return to Step 1.

Due to variable oxygen uptake rates (OUR) and changes to the anoxic timer, the overall aerobic/anoxic interval lengths typically varied between 10 – 20 minutes.

#### S1.2 Process Modeling

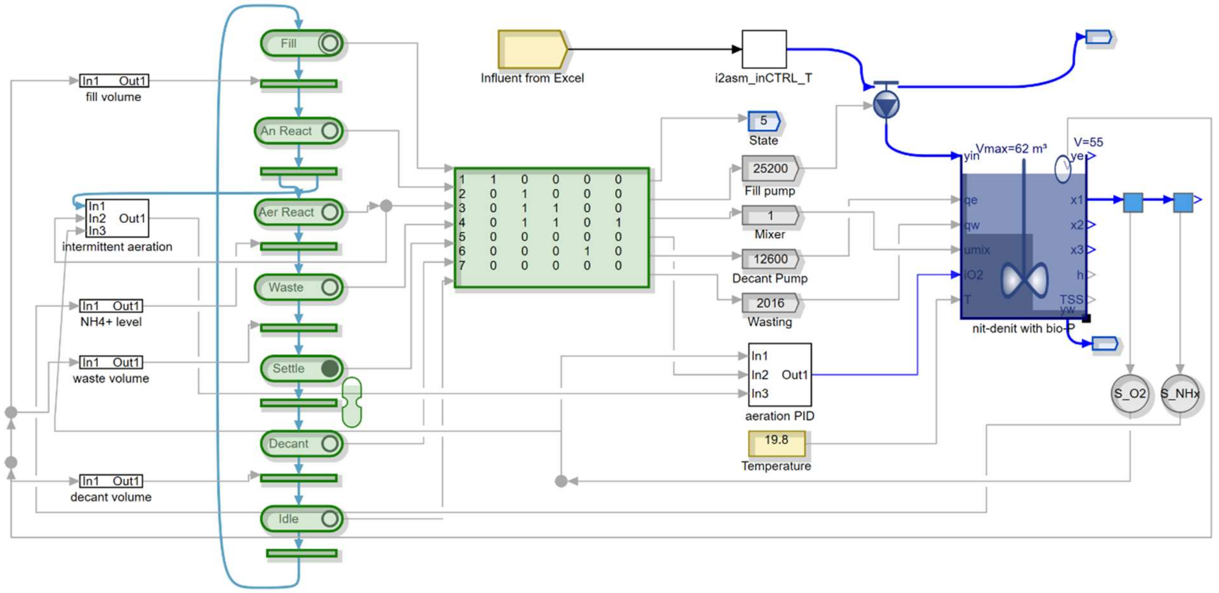


**Figure S1**. Process representation in the Simba# 3.0 software.

Modeled specific growth rates for AOO, NOO, and PAOs were quantified throughout the SBR cycles with rate equations and parameter values from the Simba# inCTRL ASM matrix. Rate equations and parameters values (at 20°C) discussed in the text are as follows:

$$\boldsymbol{net\ specific\ growth\ rate\ of\ AOO\ (d^{-1}) = \mu_{AOO}}$$

$$= \hat{\mu}_{AOO}\frac{S_{NHx}}{S_{NHx}+K_{NHx,AOO}}\frac{S_{O2}}{S_{O2}+K_{O2,AOO}}\frac{S_{PO4}}{S_{PO}+K_{PO4,ANO}}\frac{S_{ALK}}{S_{ALK}+K_{ALK,AOO}}$$

$$-\hat{b}_{AOO,O2}\frac{S_{O2}}{S_{O2}+K_{O2,AOO}}-\hat{b}_{AOO,NOx}\frac{S_{NO3}+S_{NO}}{S_{NO}+S_{NO2}+K_{NOx,ANO}}\frac{K_{O2,AOO}}{S_{O2}+K_{O2,AOO}}$$

$$-\hat{b}_{AOO,ANA}\frac{K_{NOx,ANO}}{S_{NO3}+S_{NO2}+K_{NOx,ANO}}\frac{K_{O2,AOO}}{S_{O2}+K_{O2,AOO}}$$

Where:

$$\hat{\mu}_{AOO} = maximum\ specific\ growth\ rate\ of\ AOO\ (d^{-1}) = 0.9$$

$$S_{NHx} = concentration\ of\ NH_4^+ + NH_3\ \left(\frac{mgN}{L}\right)$$

$$K_{NHx,AOO} = AOO\ half\ saturation\ coefficient\ for\ (NH_4^+ + NH_3)\left(\frac{mgN}{L}\right) = 0.7$$

$S_{O2} = concentration\ of\ dissolved\ O_2\ \left(\frac{mgO_2}{L}\right)$

$K_{O2,AOO} = AOO\ half\ saturation\ coefficient\ for\ dissolved\ O_2\ \left(\frac{mgO_2}{L}\right) = 0.25$

$S_{PO4} = concentration\ of\ PO_4^{3-}\ \left(\frac{mgP}{L}\right)$

$K_{PO4,ANO} = nitrifier\ nutrient\ half\ saturation\ coefficient\ for\ PO_4^{3-}\ \left(\frac{mgP}{L}\right) = 0.001$

$S_{ALK} = concentration\ of\ alkalinity\ \left(\frac{meq}{L}\right)$

$K_{ALK,AOO} = AOO\ half\ saturation\ coefficient\ for\ alkalinity\ \left(\frac{meq}{L}\right) = 0.5$

$\hat{b}_{AOO,O2} = maximum\ specific\ aerobic\ decay\ rate\ of\ AOO\ (d^{-1}) = 0.17$

$\hat{b}_{AOO,NOx} = maximum\ specific\ anoxic\ decay\ rate\ of\ AOO\ (d^{-1}) = 0.1$

$S_{NO3} = concentration\ of\ NO_3^-\ \left(\frac{mgN}{L}\right)$

$S_{NO2} = concentration\ of\ NO_2^-\ \left(\frac{mgN}{L}\right)$

$K_{NOx,ANO} = nitrifier\ half\ saturation\ for\ anoxic\ conditions\ \left(\frac{mgN}{L}\right) = 0.03$

$\hat{b}_{AOO,ANA} = maximum\ specific\ anaerobic\ decay\ rate\ of\ AOO\ (d^{-1}) = 0.05$

$$\boldsymbol{net\ specific\ growth\ rate\ of\ NOO\ (d^{-1}) = \mu_{NOO}}$$

$$= \hat{\mu}_{NOO}\frac{S_{NO2}}{S_{NO2}+K_{NO\ \ ,NOO}}\frac{S_{O2}}{S_{O2}+K_{O2,NOO}}\frac{S_{NHx}}{S_{NHx}+K_{NHx,ANO}}\frac{S_{PO4}}{S_{PO4}+K_{PO4,ANO}}\frac{S_{ALK}}{S_{ALK}+K_{ALK,NOO}}$$

$$-\hat{b}_{NOO,O2}\frac{S_{O2}}{S_{O2}+K_{O2,NOO}} - \hat{b}_{AOO,NOx}\frac{S_{NO3}+S_{NO2}}{S_{NO3}+S_{NO2}+K_{NOx,ANO}}\frac{K_{O2,NOO}}{S_{O2}+K_{O2,NOO}}$$

$$-\ \hat{b}_{NOO,ANA}\frac{K_{NOx,ANO}}{S_{NO3}+S_{NO2}+K_{NOx,ANO}}\frac{K_{O2,NOO}}{S_{O2}+K_{O2,NOO}}$$

Where (in addition to above):

$\hat{\mu}_{NOO} = maximum\ specific\ growth\ rate\ of\ NOO\ (d^{-1}) = 0.7$

$K_{NO2,NOO} = NOO\ half\ saturation\ coefficient\ for\ (NO_2^-)\left(\frac{mgN}{L}\right) = 0.1$

$K_{O2,NOO} = NOO\ half\ saturation\ coefficient\ for\ dissolved\ O_2\left(\frac{mgO_2}{L}\right) = 0.1$

$K_{NHx,ANO} = Nitrifier\ nutrient\ half\ saturation\ coefficient\ for\ (NH_4^+ + NH_3)\left(\frac{mgN}{L}\right) = 0.001$

$K_{ALK,NOO} = NOO\ half\ saturation\ coefficient\ for\ alkalinity\ \left(\frac{meq}{L}\right) = 0.5$

$\hat{b}_{NOO,O2} = maximum\ specific\ aerobic\ decay\ rate\ of\ NOO\ (d^{-1}) = 0.15$

$\hat{b}_{NOO,NOx} = maximum\ specific\ anoxic\ decay\ rate\ of\ NOO\ (d^{-1}) = 0.07$

$\hat{b}_{NOO,ANA} = maximum\ specific\ anaerobic\ decay\ rate\ of\ NOO\ (d^{-1}) = 0.04$

**AOO and NOO washout SRT calculation**

The modeled SRT to avoid washout for NOO was calculated by taking the inverse of average modeled $\boldsymbol{\mu_{NOO}}$ values (as shown above, calculated approximately every minute) over one cycle, i.e.:

$$\text{washout } SRT_{NOO} = \frac{1}{mean(\mu_{NOO})}$$

A similar calculation was done for AOO to affirm that modeled SRT was sufficiently high to retain AOO. The aerobic fraction of the resulting SRT for AOO and NOO was then calculated by assuming that 48% of the intermittently aerated react phase was aerobic – see Section 2.1 for details.

$$SRT_{AER} = SRT * \frac{0.48(t_{AER})}{t_{AN} + t_{AER}} = SRT * 0.399$$

Where:

$SRT_{AER} = aerobic\ SRT$

$t_{AER} = length\ of\ modeled\ intermittently\ aerated\ react\ phase\ (minutes)$

$= 222$ (variable in the actual reactor)

$t_{AN} = length\ of\ anaerobic\ react\ phase\ (minutes)$

$= 45$ (same in the actual reactor)

$\boldsymbol{specific\ growth\ rate\ of\ PAOs\ on\ PHA\ and\ O_2\ (d^{-1}) = \mu_{PAO,O2}}$

$$= \hat{\mu}_{PAO} \frac{\frac{X_{PHA}}{X_{PAO}}}{\frac{X_{PHA}}{X_{PAO}} + K_{PHA}} \frac{S_{O2}}{S_{O2} + K_{O2,OHO}} \frac{S_{NHx}}{S_{NHx} + K_{NHx,OHO}} \frac{S_{PO}}{S_{PO4} + K_{PO\ ,PAO}} \frac{S_{ALK}}{S_{ALK} + K_{ALK}}$$

Where (in addition to above):

$\hat{\mu}_{PAO} = maximum\ specific\ growth\ rate\ of\ PAOs\ (d^{-1}) = 0.95$

$X_{PHA} = concentration\ of\ polyhydroxyalkanoates - PHAs\ \left(\frac{mgCOD}{L}\right)$

$X_{PAO} = concentration\ of\ PAOs\ \left(\frac{mgCOD}{L}\right)$

$K_{PHA} = half\ saturation\ coefficient\ for\ PHA\ \left(\frac{mgCOD}{L}\right) = 0.1$

$K_{O2,OHO} = OHO\ and\ PAO\ half\ saturation\ coefficient\ for\ dissolved\ O_2\ \left(\frac{mgO_2}{L}\right) = 0.05$

$K_{NHx,OHO} = OHO\ and\ PAO\ nutrient\ half\ saturation\ coefficient\ for\ (NH_4^+ + NH_3)\left(\frac{mgN}{L}\right) = 0.001$

$K_{PO4,PAO} = PAO\ half\ saturation\ coefficient\ for\ PO_4^{3-}\ \left(\frac{mgP}{L}\right) = 0.15$

$K_{ALK} = PAO\ half\ saturation\ coefficient\ for\ alkalinity\ \left(\frac{meq}{L}\right) = 0.1$

$\boldsymbol{specific\ growth\ rate\ of\ PAOs\ on\ PHA\ and\ NO_2^-\ (d^{-1}) = \mu_{PAO,NO2}}$

$$= \hat{\mu}_{PAO}\,\eta_{anox,PAO}\frac{\frac{X_{PHA}}{X_{PAO}}}{\frac{X_{PHA}}{X_{PAO}} + K_{PHA}}\frac{S_{NO}}{S_{NO2} + K_{NO\ \ ,OHO}}\frac{K_{O2,OHO}}{S_{O2} + K_{O2,OHO}}\frac{S_{NHx}}{S_{NHx} + K_{NHx,OHO}}$$

$$\frac{S_{PO}}{S_{PO4} + K_{PO4,PAO}}\frac{S_{ALK}}{S_{ALK} + K_{ALK}}$$

Where (in addition to above):

$\eta_{anox,PAO} = PAO\ anoxic\ growth\ factor = 0.33$

$K_{NO\ \ ,OHO} = OHO\ and\ PAO\ half\ saturation\ coefficient\ for\ NO_2^-\left(\frac{mgN}{L}\right) = 0.05$

$\boldsymbol{specific\ growth\ rate\ of\ PAOs\ on\ PHA\ and\ NO_3^-\ (d^{-1}) = \mu_{PAO,NO3}}$

$$= \hat{\mu}_{PAO}\,\eta_{anox,PAO}\frac{\frac{X_{PHA}}{X_{PAO}}}{\frac{X_{PHA}}{X_{PAO}} + K_{PHA}}\frac{S_{NO}}{S_{NO3} + K_{NO3,OHO}}\frac{K_{NO\ \ ,OHO}}{S_{NO2} + K_{NO\ \ ,OHO}}\frac{K_{O2,OHO}}{S_{O2} + K_{O2,OHO}}$$

$$\frac{S_{NHx}}{S_{NHx} + K_{NHx,OHO}}\frac{S_{PO4}}{S_{PO4} + K_{PO\ \ ,PAO}}\frac{S_{ALK}}{S_{ALK} + K_{ALK}}$$

Where (in addition to above):

$K_{NO\ \ ,OHO} = OHO\ and\ PAO\ half\ saturation\ coefficient\ for\ NO_3^-\left(\frac{mgN}{L}\right) = 0.1$

**S1.3. Solids Retention Time (SRT) Control**

SRT was controlled via timed mixed liquor wasting after the aerated react period and before settling. A maximum wasting pump time was set on the PLC, and the actual pumping time for each cycle varied depending on the length of the aerated react phase. For example, if the maximum wasting pump time was set to 1 minute, the maximum aeration time was set to 300 minutes, and the actual aeration time for a given cycle was 150 minutes, the actual pumping time would be $1\ minute \times \frac{150\ minutes}{300\ minutes} = 0.5\ minutes$. Because the aeration time varied on a cycle-by-cycle basis according to the influent strength and the target effluent $NH_4^+$ level, the dynamic SRT value was calculated for each individual cycle, as adapted from Laureni et al. (2019) and Takács et al., (2008). SRT for each cycle was calculated according to the equation below (Laureni et al., 2019).

$$SRT_{t+\Delta t} = SRT_t\left(1 - \frac{X_E V_E + X_R V_W}{X_R V_R}\right) + \Delta t$$

Where:

| | |
|---|---|
| $SRT_{t+\Delta t}$ | = Solids retention time of cycle under analysis (days) |
| $SRT_t$ | = Solids retention time of previous cycle (days) |
| $V_R$ | = Volume of reactor (L) |
| $X_E$ | = Effluent VSS concentration for the cycle under analysis (mg/L) |
| $V_E$ | = Effluent volume for the cycle under analysis (L) |
| $X_R$ | = Reactor MLVSS concentration for the cycle under analysis (mg/L) |
| $V_W$ | = Mixed liquor wasting volume for the cycle under analysis (L) |
| $\Delta t$ | = React time of the cycle under analysis, not including settling and decant (days) |

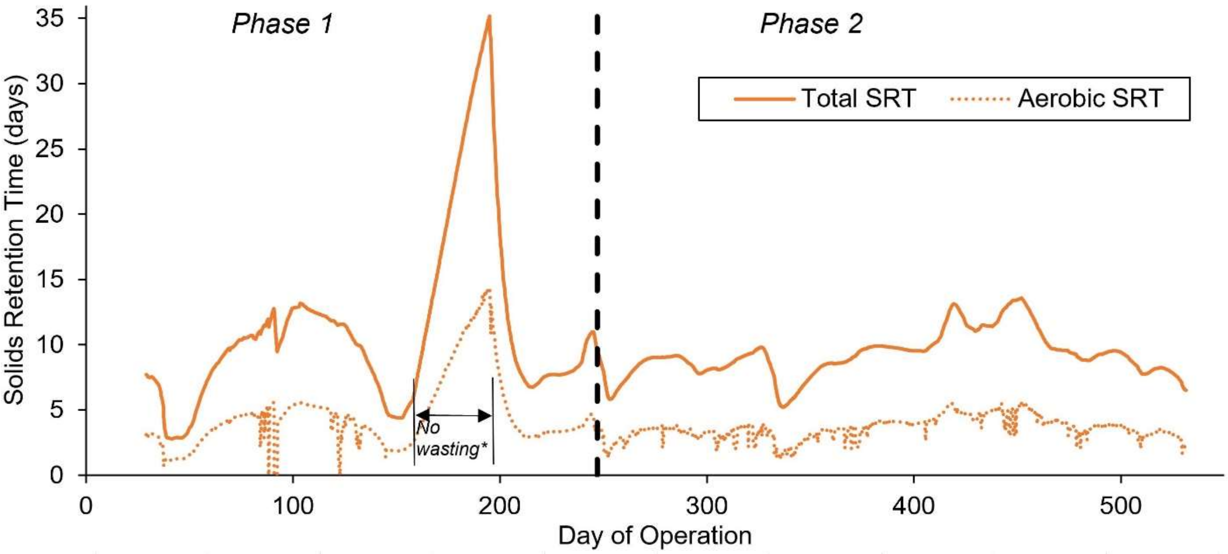


**Figure S2**. Total and aerobic dynamic SRT over time in the SBR. The average total and aerobic SRT during Phase 1 was 11 ± 7 and 4.5 ± 3.0 days, and the average total and aerobic SRT during Phase 2 was 9.2 ± 1.8 and 3.6 ± 0.9 days, respectively. *Mixed liquor wasting was suspended from days 158 – 195 to recover AOO activity.

**S1.4. 16S rRNA Gene Amplicon Sequencing**

16S rRNA gene amplicon library preparations were performed using a two-step PCR protocol using the Fluidigm Biomark: Multiplex PCR Strategy as previously described (Griffin and Wells, 2017). In the first round of PCR, each 20 uL reaction contained 10 µL of FailSafe PCR 2X PreMix F (Epicentre, Madison, WI), 0.63 units of Expand High Fidelity PCR Taq Enzyme (Sigma-Aldrich, St. Louis, MO), 0.4 µM of forward primer and reverse primer modified with Fluidigm common sequences at the 5' end of each primer, 1 µL of gDNA (approximately 100 ng) and the remaining volume molecular biology grade water. The V4-V5 region of the 16S rRNA gene was amplified in duplicate from 10 samples collected over the course of reactor operation using the 515F-Y (5'-GTGYCAGCMGCCGCGGTAA-3') and 926R (5'-CCGYCAATTYMTTTRAGTTT-3') (Parada et al., 2016) primer set. Thermocycling conditions for the 515F-Y/926R primer set were 95°C for 5 minutes, then 28 cycles of 95°C for 30 seconds, 50°C for 45 seconds, and 68°C for 30 seconds, followed by a final extension of 68°C for 5 minutes. Specificity of amplification was checked for all samples via agarose gel electrophoresis.

Samples were then barcoded by sample via a second stage PCR amplification using Access Array Barcodes (Fluidigm, South San Francisco, CA) (Griffin and Wells, 2017). Each 20 uL PCR reaction consisted of 10 µL of FailSafe PCR 2X PreMix F, 0.63 units of Expand High Fidelity PCR Taq Enzyme, 2 µL of template from the first round of PCR, 4 µL of sample-specific barcode primers and the remaining volume molecular biology grade water. The conditions for the second round of PCR were 95°C for 5 minutes, then 8 cycles of 95°C for 30 seconds, 60°C for 30 seconds, and 68°C for 30 seconds. Agarose gel electrophoresis was run again after the second round of PCR to verify correct amplification. Sequencing was performed on an Illumina Miseq sequencer (Illumina, San Diego, CA) using Illumina V2 (2x250 paired end) chemistry.

For amplicon sequence analysis, sequence quality control was performed through DADA2 (Callahan et al., 2016) integrated in QIIME2 version qiime2-2018.8 (Bolyen et al., 2018), which included quality-score-based sequence truncation, primer trimming, merging of paired-end reads, and removal of chimeras. Taxonomy was assigned to each individual sequence variation using the Silva database, release 132.

**S1.5 qPCR supermix and reaction conditions**

Bio-Rad iQ SYBR Green Supermix (Bio-Rad, Hercules, CA, USA) containing 50 U/ml iTaq DNA polymerase, 0.4 mM dNTPs, 100 mM KCl, 40 mM Tris-HCl, 6 mM MgCl2, 20 mM fluorescein, and stabilizers was used for two qPCR assays. Target genes included ammonia oxidizing bacterial *amoA* via the *amoA*-1F and *amoA*-2R primer set (Rotthauwe et al., 1997) and total bacterial (universal) 16S rRNA genes via the Eub519/Univ907 primer set (Burgmann et al., 2011). The final volume of the reaction mix for each PCR and qPCR reaction was 20 µl, in which the DNA template was ~1 ng, and the primer concentrations were 0.2 μM. All assays were performed in triplicate. For each assay, triplicate standard series were generated by tenfold serial dilutions ($10^2$-$10^8$ gene copies/μl).

### S2. Process Modeling Reproduces Key Elements of Process Performance

Agreement between the process model and our experimental results suggest that the trends in N and P removal from mainstream wastewater that we observed are likely generally applicable to other locations. By closely modeling the influent (primary effluent), reactor control, aeration control and SRT (model SRT 9.5 days, reactor SRT 9.2 ± 1.8 days) from Phase 2, the resulting model performance closely matched that of the reactor (Figure 5): modeled HRT was 7.2 hours (reactor HRT 6.8 ± 2.8 hours), modeled VSS was 1,245 mg/L (reactor VSS 1,344 ± 226 mg/L) and Figure 2, Figure 4, and Table 3 demonstrate that both in-cycle nutrient dynamics and effluent concentrations were well-matched between the model and reactor performance. Importantly, this was done via a commercially available wastewater process modeling software without modification to the inCTRL ASM matrix.

### S3. Supporting Table and Figures

**Table S1.** Influent (primary effluent) COD fractionation and COD-to-nutrient ratios.

| | Primary Effluent | As percent of total COD |
|---|---|---|
| Total COD (mgCOD/L)[a] | 164.4 ± 46.2 | --- |
| Particulate COD (mgCOD/L) | 61.7 ± 23.8 | 37% |
| Colloidal COD (mgCOD/L) | 28.6 ± 18.1 | 17% |
| Soluble COD not including VFA (mgCOD/L) | 56.4 ± 19.4 | 34% |
| VFA (mgCOD/L) | 18.8 ± 8.9 | 11% |
| COD:TP[b] (gCOD/gP) | 67:1 | --- |
| COD:TKN[b] (gCOD/gN) | 8.3:1 | --- |

[a]Primary effluent COD fractionation was performed weekly from days 114 - 515 (n = 50).

[b]COD:Nutrient ratios are taken from average of all samples from days 27 - 519 (n = 192).

**Table S2.** $N_2O$ emissions test results for 8 cycles during Phase 2.

| Day of cycle tested | $N_2O$ emitted/ influent TKN | $N_2O$ emitted/ TIN removed | influent TKN (mgN/L) | influent COD (mg/L) | COD/ TKN | Effluent $NO_2^-$ (mgN/L) | Average temp (°C) |
|---|---|---|---|---|---|---|---|
| 414 | 3.8% | 11.4% | 23 | 206 | 9 | 2.9 | 20.5 |
| 426 | 6.2% | 12.0% | 20 | 204 | 10 | 2.7 | 20.3 |
| 428 | 1.0% | 2.3% | 12 | 140 | 12 | 1.2 | 20.5 |
| 475 | 1.0% | 2.6% | 13 | 64 | 5 | 2.0 | 20.4 |
| 489 | 2.2% | 4.3% | 19 | 183 | 10 | 2.4 | 20.3 |
| 503 | 0.2% | 0.2% | 14 | 160 | 11 | 0.4 | 19.4 |
| 517 | 0.8% | 1.6% | 21 | 147 | 7 | 1.9 | 19.4 |
| 531 | 1.56% | 7.36% | 13 | 144 | 11 | 2.1 | 19.4 |

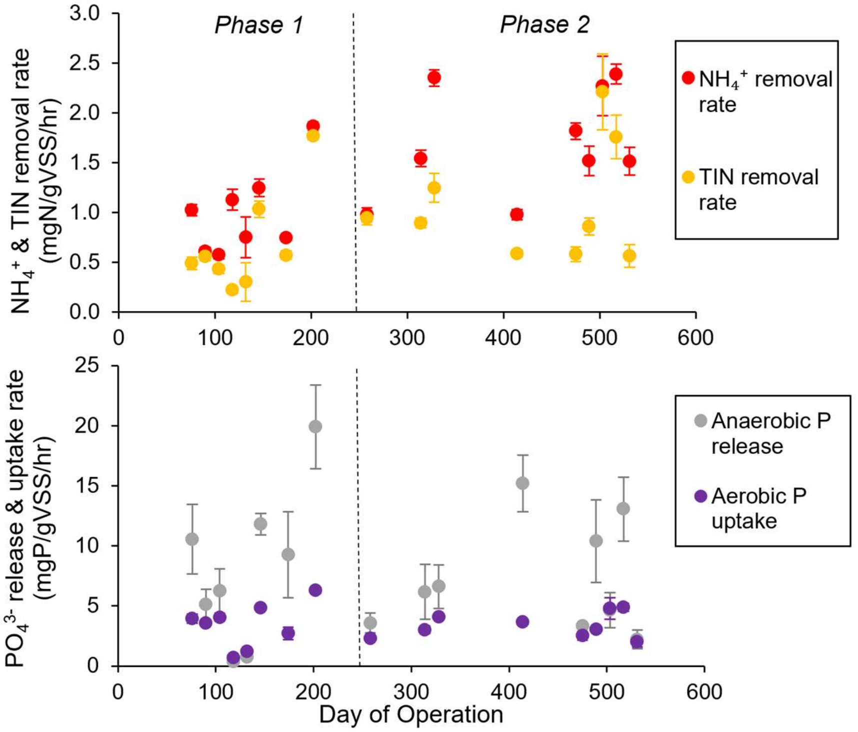


**Figure S3.** In-cycle N and P removal rates from least-squares regression of the linear portions of in-cycle grab samples for $NH_4^+$, $NO_2^-$, $NO_3^-$, and $PO_4^{3-}$. Error bars represent standard errors of the slopes.

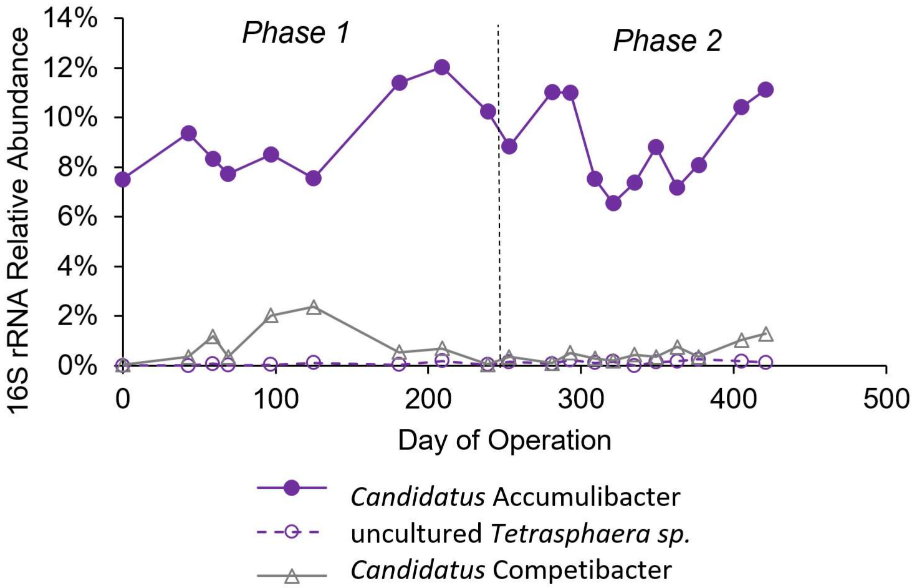


**Figure S4.** Relative *Accumulibacter*, *Tetrasphaera*, and *Competibacter* abundance through the first 421 days of reactor operation according to 16S rRNA gene sequencing. Day "0" represents the inoculum, which was sampled before reactor operation began.

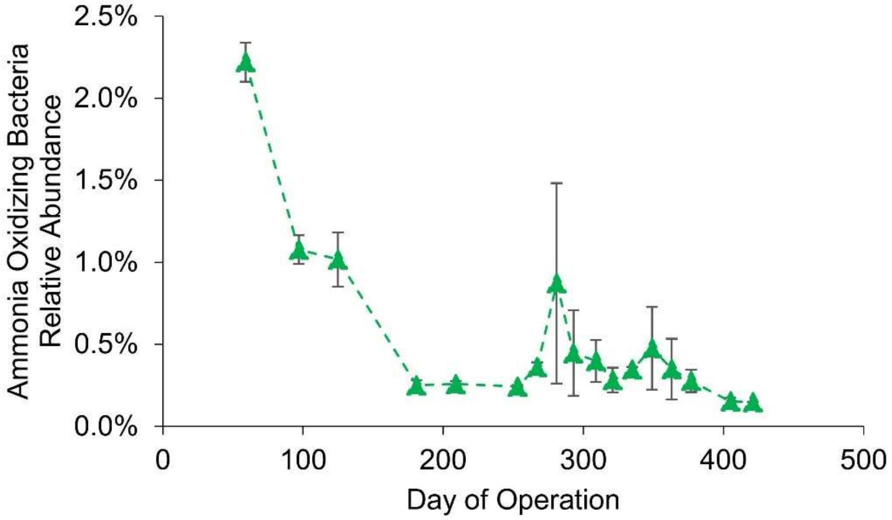


**Figure S5.** Ammonia oxidizing bacterial *amoA* gene abundance normalized to total bacterial 16S rRNA genes through the first 421 days of reactor operation according to qPCR.

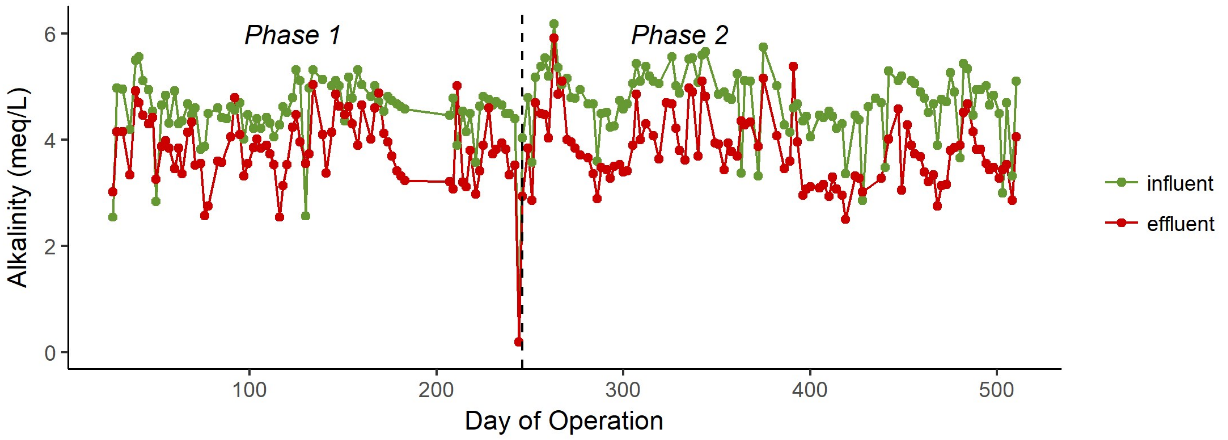


**Figure S6.** Reactor influent and effluent alkalinity concentrations from composite sampling

.

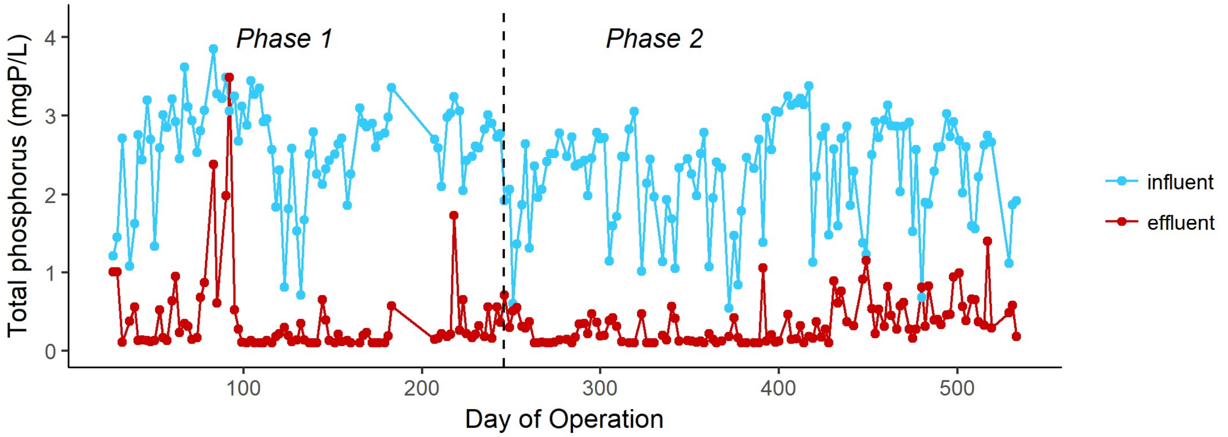


**Figure S7.** Reactor influent and effluent total phosphorus concentrations from composite sampling.

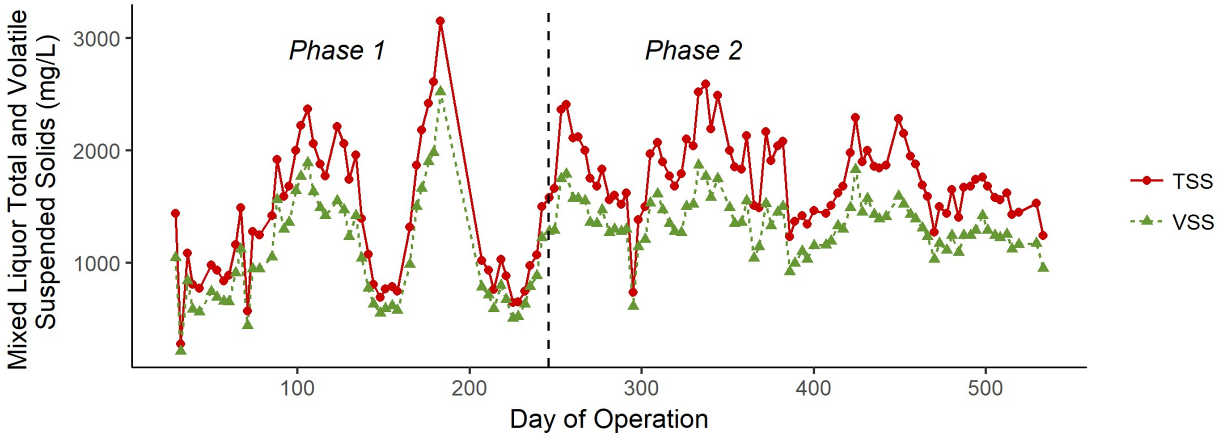


**Figure S8.** Reactor mixed liquor TSS and VSS concentrations.

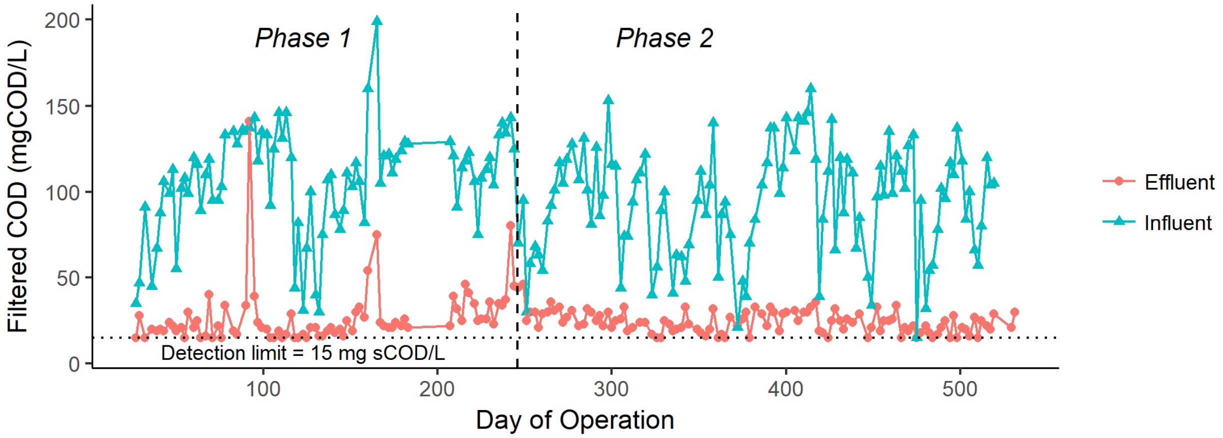


**Figure S9.** Reactor influent and effluent filtered COD from composite sampling. Samples were filtered through a 1.2 μm pore size membrane.

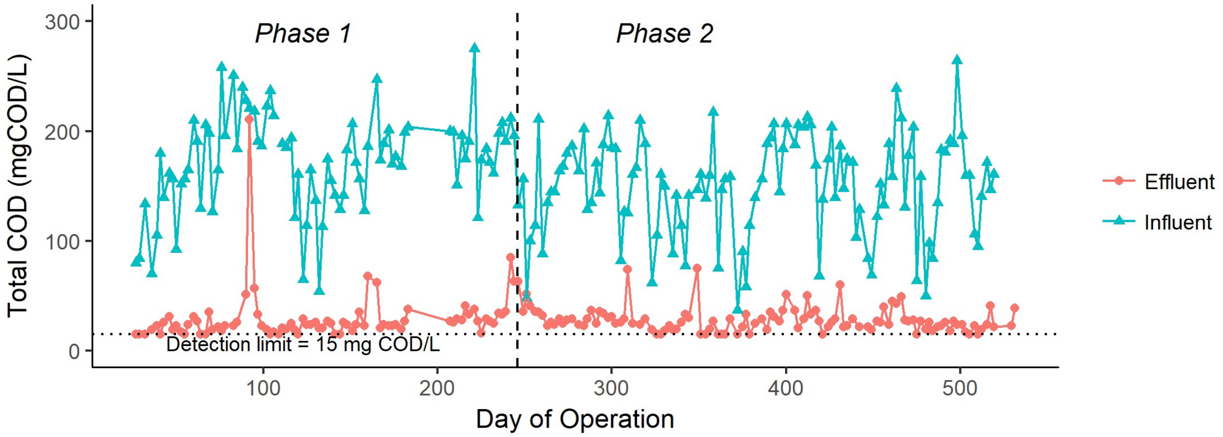


**Figure S10.** Reactor influent and effluent total COD concentrations from composite sampling.

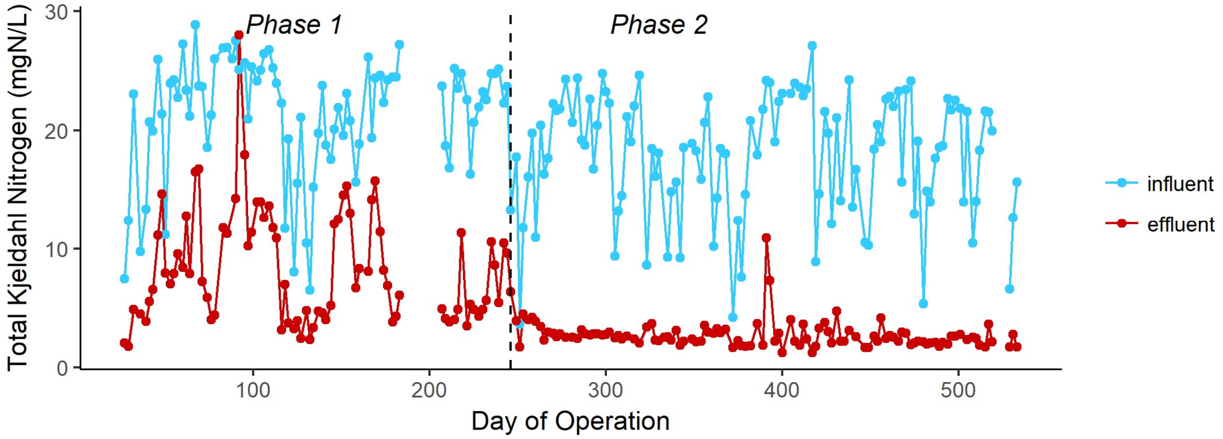


**Figure S11.** Reactor influent and effluent TKN concentrations from composite sampling.

### S3. References for Supporting Information


Bolyen, E., Rideout, J.R., Dillon, M.R., Bokulich, N.A., Abnet, C., Al-Ghalith, G.A., Alexander, H., Alm, E.J., Arumugam, M., Asnicar, F., Bai, Y., Bisanz, J.E., Bittinger, K., Brejnrod, A., Brislawn, C.J., Brown, C.T., Callahan, B.J., Caraballo-Rodríguez, A.M., Chase, J., Cope, E., Da Silva, R., Dorrestein, P.C., Douglas, G.M., Durall, D.M., Duvallet, C., Edwardson, C.F., Ernst, M., Estaki, M., Fouquier, J., Gauglitz, J.M., Gibson, D.L., Gonzalez, A., Gorlick, K., Guo, J., Hillmann, B., Holmes, S., Holste, H., Huttenhower, C., Huttley, G., Janssen, S., Jarmusch, A.K., Jiang, L., Kaehler, B., Kang, K.B., Keefe, C.R., Keim, P., Kelley, S.T., Knights, D., Koester, I., Kosciolek, T., Kreps, J., Langille, M.G., Lee, J., Ley, R., Liu, Y.-X., Loftfield, E., Lozupone, C., Maher, M., Marotz, C., Martin, B., McDonald, D., McIver, L.J., Melnik, A.V., Metcalf, J.L., Morgan, S.C., Morton, J., Naimey, A.T., Navas-Molina, J.A., Nothias, L.F., Orchanian, S.B., Pearson, T., Peoples, S.L., Petras, D., Preuss, M.L., Pruesse, E., Rasmussen, L.B., Rivers, A., Robeson, II, M.S., Rosenthal, P., Segata, N., Shaffer, M., Shiffer, A., Sinha, R., Song, S.J., Spear, J.R., Swafford, A.D., Thompson, L.R., Torres, P.J., Trinh, P., Tripathi, A., Turnbaugh, P.J., Ul-Hasan, S., van der Hooft, J.J., Vargas, F., Vázquez-Baeza, Y., Vogtmann, E., von Hippel, M., Walters, W., Wan, Y., Wang, M., Warren, J., Weber, K.C., Williamson, C.H., Willis, A.D., Xu, Z.Z., Zaneveld, J.R., Zhang, Y., Knight, R., Caporaso, J.G., 2018. QIIME 2: Reproducible, interactive, scalable, and extensible microbiome data science. PeerJ. https://doi.org/10.7287/peerj.preprints.27295v1

Burgmann, H., Jenni, S., Vazquez, F., Udert, K.M., 2011. Regime Shift and Microbial Dynamics in a Sequencing Batch Reactor for Nitrification and Anammox Treatment of Urine. Appl. Environ. Microbiol. 77, 5897–5907. https://doi.org/10.1128/AEM.02986-10

Callahan, B.J., McMurdie, P.J., Rosen, M.J., Han, A.W., Johnson, A.J.A., Holmes, S.P., 2016. DADA2: High-resolution sample inference from Illumina amplicon data. Nat. Methods 13, 581–583. https://doi.org/10.1038/nmeth.3869

Griffin, J.S., Wells, G.F., 2017. Regional synchrony in full-scale activated sludge bioreactors due to deterministic microbial community assembly. ISME J. 11, 500–511. https://doi.org/10.1038/ismej.2016.121

Laureni, M., Weissbrodt, D.G., Villez, K., Robin, O., de Jonge, N., Rosenthal, A., Wells, G., Nielsen, J.L., Morgenroth, E., Joss, A., 2019. Biomass segregation between biofilm and flocs improves the control of nitrite-oxidizing bacteria in mainstream partial nitritation and anammox processes. Water Res. 154, 104–116. https://doi.org/10.1016/j.watres.2018.12.051

Parada, A.E., Needham, D.M., Fuhrman, J.A., 2016. Every base matters: assessing small subunit rRNA primers for marine microbiomes with mock communities, time series and global field samples: Primers for marine microbiome studies. Environ. Microbiol. 18, 1403–1414. https://doi.org/10.1111/1462-2920.13023

Rotthauwe, J.-H., Witzel, K.-P., Liesack, W., 1997. The ammonia monooxygenase structural gene amoA as a functional marker: molecular fine-scale analysis of natural ammonia-oxidizing populations. Appl. Environ. Microbiol. 63, 4704–4712.

Takács, I., Stricker, A.-E., Achleitner, S., Barrie, A., Rauch, W., Murthy, S., 2008. Do You Know Your Sludge Age? Proc. Water Environ. Fed. 2008, 3639–3655. https://doi.org/10.2175/193864708788733486
